# Anaerobic degradation of syringic acid by an adapted strain of *Rhodopseudomonas palustris*

**DOI:** 10.1101/740985

**Authors:** J. Zachary Oshlag, Yanjun Ma, Kaitlin Morse, Brian T. Burger, Rachelle A. Lemke, Steven D. Karlen, Kevin S. Myers, Timothy J. Donohue, Daniel R. Noguera

## Abstract

While lignin represents a major fraction of the carbon in plant biomass, biological strategies to convert the components of this heterogenous polymer into products of industrial and biotechnological value are lacking. Syringic acid (3,5-dimethoxy-4-hydroxybenzoic acid) is a byproduct of lignin degradation, appearing in lignocellulosic hydrolysates, deconstructed lignin streams, and other agricultural products. *Rhodopseudomonas palustris* CGA009 is a known degrader of phenolic compounds under photoheterotrophic conditions, via the benzoyl-CoA degradation (BAD) pathway. However, *R. palustris* CGA009 is reported to be unable to metabolize *meta*-methoxylated phenolics such as syringic acid. We isolated a strain of *R. palustris* (strain SA008.1.07), adapted from CGA009, which can grow on syringic acid under photoheterotrophic conditions, utilizing it as a sole source of organic carbon and reducing power. An SA008.1.07 mutant with an inactive benzoyl-CoA reductase structural gene was able to grow on syringic acid, demonstrating that the metabolism of this aromatic compound is not through the BAD pathway. Comparative gene expression analyses of SA008.1.07 implicated the involvement of products of the *vanARB* operon (*rpa3619-rpa3621*), which has been described as catalyzing aerobic aromatic ring demethylation in other bacteria, in anaerobic syringic acid degradation. In addition, experiments with a *vanARB* deletion mutant demonstrated the involvement of the *vanARB* operon in anaerobic syringic acid degradation. These observations provide new insights into the anaerobic degradation of *meta*-methoxylated and other aromatics by *R. palustris*.

**IMPORTANCE:** Lignin is the most abundant aromatic polymer on Earth and a resource that could eventually substitute for fossil fuels as a source of aromatic compounds for industrial and biotechnological applications. Engineering microorganisms for production of aromatic-based biochemicals requires detailed knowledge of metabolic pathways for the degradation of aromatics that are present in lignin. Our isolation and analysis of a *Rhodopseudomonas palustris* strain capable of syringic acid degradation reveals a previously unknown metabolic route for aromatic degradation in *R. palustris*. This study highlights several key features of this pathway and sets the stage for a more complete understanding of the microbial metabolic repertoire to metabolize aromatic compounds from lignin and other renewable sources.

## INTRODUCTION

As one of the major biopolymers present in plant tissues, lignin has the potential to serve as a renewable source of carbon for the bio-based production of compounds that are currently derived from petroleum. Unfortunately, the ability to derive chemicals of commercial, chemical, or medicinal value from lignin is limited by information needed to improve the biological conversion of the aromatics in lignin into valuable products. We are interested in improving our understanding of how bacteria metabolize the aromatic building blocks in lignin and using this information to develop strategies that allow the conversion of this major component of plant cell walls into valuable products.

Syringic acid, along with other *meta*-methoxy substituted phenolic compounds are plant-derived aromatics that present both a hindrance and a potential source of value to the chemical, fuel, and biotechnology industries (1–3). Originating from the guaiacyl (coniferyl alcohol) and syringyl (sinapyl alcohol) phenylpropanoids that are polymerized into lignin during secondary cell wall formation (1), *meta*-methoxylated aromatics are frequently present in products generated from deconstructed biomass (4). While present at low concentrations in sugar-rich lignocellulosic hydrolysates, these methoxylated aromatics can nonetheless induce stress responses (2, 5) and cause toxicity (6, 7) in non-aromatic degrading microbes, leading to a decrease in both microbial growth and biofuel yield during fermentation (8, 9). Further, these phenolics are present at much higher concentrations in solubilized lignin streams produced with emerging technologies (10–13). Incorporation of *meta*-methoxylated aromatics into the metabolism of an appropriate, genetically tractable microorganism could provide a promising and efficient route for monolignol valorization through the identification and optimization of the biochemical pathways involved.

To expand the ability of microbes to metabolize syringic acid and related plant-derived aromatic compounds, we are studying *Rhodopseudomonas palustris*, a metabolically versatile, well-characterized, and genetically tractable purple non-sulfur α-proteobacterium (14–16) that has a proven and well understood ability to utilize aromatic monomers (17, 18). Under anaerobic conditions, *R. palustris* uses the benzoyl-CoA degradation (BAD) pathway to cleave the aromatic ring of mono-aromatic compounds after activation of the molecule via coenzyme-A ligation (19). The diversity of aromatic compounds that *R. palustris* can degrade depends on the existence of accessory pathways that transform aromatic monomers to the common BAD pathway intermediates benzoyl-CoA or 4-hydroxybenzoyl-CoA (20, 21). In addition, previous studies have shown that growth of *R. palustris* in lignocellulosic hydrolysates that contain a mixture of plant-derived organic compounds allows for the degradation of aromatic monomers that do not support growth when supplied as the sole carbon source in defined media (21).

Here we describe studies aimed at understanding the metabolism of syringic acid by an adapted *R. palustris* strain. By supplying syringic acid to a series of successive cultures, we isolated a strain of *R. palustris* capable of utilizing this *meta*-methoxylated aromatic as the sole source of organic carbon. We analyze the degradation of syringic acid by this adapted isolate, *R. palustris* SA008.1.07, in defined laboratory media to provide insight into the mechanisms involved in the degradation of this aromatic monomer.

## MATERIALS AND METHODS

### Media

All *R. palustris* strains were grown in PM medium (22), brought to pH 7 with sodium hydroxide, and sterilized by filtration. PM media with different organic carbon sources were prepared; PM-AcY contained 20 mM sodium acetate and 0.1% yeast extract, PM-succinate contained 10 mM succinic acid, PM-aromatic was made with 3 to 3.5 mM aromatic compounds (unless otherwise indicated), and supplemented with 30 mM sodium bicarbonate. *Escherichia coli* strains were grown on LB medium (23). Molecular genetics-grade agar (Fisher Scientific, Fair Lawn, NJ) was added to media at 1.5% to solidify, where noted. When necessary, the following reagents were used for cloning, selection, and propagation of modified strains: sucrose 10% (w/v), kanamycin (Kn) 50 µg/mL, ampicillin 25 µg/mL, gentamycin 20 µg/mL. All chemicals for media preparation were obtained from Fisher Scientific (Hampton, NH) or Sigma-Aldrich (St. Louis, MO) at purities suitable for molecular biology.

### Strains and Plasmids

The *E. coli* and *R. palustris* strains and plasmids used in this study are summarized in Table 1.

**Table 1.**
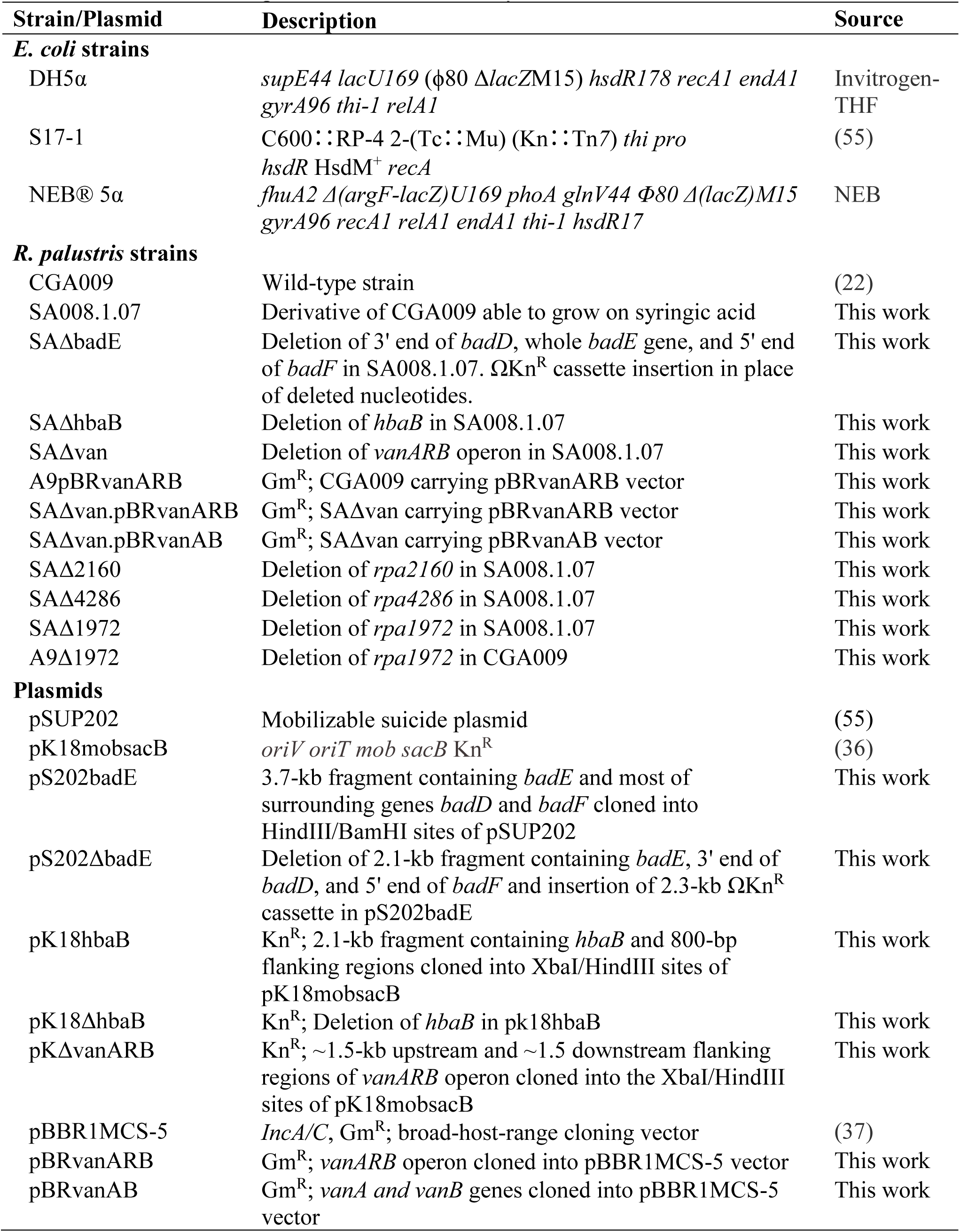
List of strains and plasmids used in this study

### Growth Conditions

To culture *R. palustris,* cells were streaked from glycerol freezer stocks onto PM-AcY-agar plates and incubated aerobically at 30 °C to obtain single colonies. A colony was transferred to 25 ml PM-AcY liquid media and grown aerobically at 30 °C. Aliquots (about 170 μl) from this aerobic culture were added to clear glass culture tubes (16×125 mm) containing PM-succinate, which were completely filled to the brim with media, sealed with a rubber septum, and incubated. Since the growing culture rapidly exhausts any oxygen available in the medium, this culturing technique has been demonstrated to efficiently create anaerobic culturing conditions in liquid media (24). Photoheterotrophic growth was maintained at 30 °C under illumination by incandescent tungsten lamps at ∼10 W/m^2^ and kept well mixed by a micro magnetic stir bar (3×10mm). These photoheterotrophic PM-succinate cultures were used as inocula for photoheterotrophic experiments with the aromatic substrates, which were prepared following the procedure described above to generate anaerobic conditions in liquid media. *R. palustris* growth in liquid cultures was monitored using a Klett-Summerson photoelectric colorimeter (Klett MFG Co., New York, NY). Photoheterotrophic growth on solid media was achieved by placing plates in a sealed canister containing a GasPak^TM^ Ez Anaerobe Container System (BD Biosciences, Franklin Lakes, NJ), which was placed under constant illumination and rotated daily.

### Analytical tests

For chemical analysis, samples were taken periodically by aseptically piercing a rubber septum and withdrawing 200 µL of liquid culture. Following sampling, the headspace of cultures was flushed with argon gas. Samples were passed through 0.22 µm PVDF membranes (Merck, KGaA, Darmstadt, Germany) to separate cells from media, and the filtrates were frozen at −80 °C until analysis.

Aromatic compounds were quantified by high performance liquid chromatography (HPLC) using an LC-10AT_VP_ solvent delivery module HPLC system (Shimadzu, Kyoto, Japan) with an SPD-M10A_VP_ diode array detector (Shimadzu, Kyoto, Japan). Samples were prepared as described elsewhere (25). Aromatic compounds were separated using a C18-reversed stationary phase column and an isocratic aqueous mobile phase of methanol (30% [wt/vol]), acetonitrile (6% [wt/vol]), and 5 mM formic acid in water (64% [wt/vol]) at a flow rate of 0.8 ml min^−1^ (25). Aromatics and metabolic by-products were quantified using standard curves and UV absorbance. Standard curves were prepared using commercially purchased compounds (Sigma-Aldrich, St. Louis, MO) dissolved in dimethyl sulfoxide (DMSO).

Liquid chromatography tandem-mass spectrometry (LC-MS/MS) was used for identification of extracellular metabolic by-products, using a chromatography separation system similar to the one described above. Mass spectra were analyzed with a Thermo Q-exactive mass spectrometer (Thermo Scientific, Waltham, MA). Standards were directly infused into the mass spectrometer. Spectra were acquired in positive ionization mode with an MS/MS resolution of 17,500, isolation width 2.0 Da, and normalized collision energy 30%.

Nuclear Magnetic Resonance (NMR) was also used for identification of some metabolic by-products. For these tests, three consecutive 100-mL ethyl acetate (EtOAc) extractions were performed on 500 mL of spent medium at pH 6.5-7.0. The pH of the aqueous fraction was then lowered to ∼1 using 1 M hydrochloric acid. Additional organic compounds were extracted from this acidified aqueous fraction using three consecutive 100-mL dichloromethane (DCM) extractions. Both extractants were independently washed three times with saturated sodium bicarbonate (50 mL/extraction), then twice with saturated sodium chloride (50 mL/extraction), and then dried with sodium sulfate. Samples were filtered and the solvent was evaporated. NMR spectra of the extracted compounds were collected in acetone-*d_6_* on a Bruker AVANCE 500 MHz spectrometer (Billerica, MA, USA) fitted with a cryogenically-cooled 5-mm QCI (1H/31P/13C/15N) gradient probe with inverse geometry (proton coils closest to the sample). Spectra were compared to high purity standards from Sigma-Aldrich (St. Louis, MO).

Chemical Oxygen Demand (COD) was used to quantify soluble organic compounds and biomass (25), with measurements on both filtered and unfiltered samples. The theoretical COD values for various carbon sources used in this study are as follows (in mg of COD/mmol of substrate): benzoic acid 240, 4-hydroxybenzoic acid (4-HBA) 224; syringic acid 288.

### Transcriptomic analysis (RNA-seq)

For transcriptomic analyses, *R. palustris* SA008.1.07 cultures were photoheterotrophically grown on PM-4-HBA, PM-syringic acid, or PM-succinate by bubbling with 95% N_2_ and 5% CO_2_ under constant illumination at 30°C to mid-log phase when RNA was harvested (26). For each sample, rRNA was reduced (Ribo-Zero kit, Illumina), and a strand-specific library was prepared (TruSeq Stranded Total RNA Sample Prep Kit, Illumina). RNA from cultures grown on PM-4-HBA and PM-syringic acid was processed and sequenced at the University of Wisconsin-Madison Biotechnology Center (Illumina HiSeq2500, 1×100 bp, single end). RNA from cultures grown on PM-succinate was processed and sequenced at the U.S. Department of Energy Joint Genome Institute (Illumina NextSeq, 2×151 bp, paired end). Three biological replicates were analyzed per growth condition. The paired-end FASTQ files were split into read 1 (R1) and read 2 (R2) files and R1 files were retained for further analysis as the other data contained only single-end reads. All FASTQ files were processed through the same pipeline. Reads were trimmed using Trimmomatic version 0.3 (27) with the default settings except for a HEADCROP of 5, LEADING of 3, TRAILING of 3, SLIDINGWINDOW of 3:30, and MINLEN of 36. After trimming, the reads were aligned to the *R. palustris* CGA009 genome sequence (GenBank assembly accession GCA_000195775.1) using Bowtie2 version 2.2.2 (28) with default settings except the number of mismatches allowed was set to 1. Aligned reads were mapped to gene locations using HTSeq version 0.6.0 (29) using default settings except for the “reverse” strandedness argument was used. DESeq2 version 1.22.2 (30) was used to identify significantly differentially expressed genes from pairwise analyses, using a Benjamini and Hochberg (31) false discovery rate (FDR) less than 0.05 as a significance threshold and/or a fold change greater than two. Raw sequencing reads were normalized using the reads per kilobase per million mapped reads (RPKM). A full list of gene transcripts normalized by RPKM is shown in Table S1. The accession number for the RNA-sea data in the Gene Expression Omnibus (GEO) database is GSE135630.

### Genome sequencing

Genomic DNA of CGA009, SA008.1.07 and 16 other adapted strains which were capable of anaerobic degradation of syringic acid was isolated and purified (32). Genome sequencing was performed and analyzed by the U.S. Department of Energy Joint Genome Institute on an Illumina NovaSeq (2 x151 bp). The resulting DNA reads were aligned to the *R. palustris* CGA009 genome (NC005296) using the short read alignment tool BWA (33). SNPs and small INDELs were called using samtools mpileup and bcftools then filtered using vcfutils.pl from the samtools package (34). The NCBI accession numbers for sequences are PRJNA520130-PRJNA520144, PRJNA537839, and PRJNA537840.

### DNA manipulation

Purification of PCR products was achieved using the QIAquick PCR purification kit (Qiagen, Hilden, GER) and PCR products were extracted and purified from agarose gels using the Zymoclean Gel DNA Recovery Kit (Zymo Research, Irvine, CA). The Zyppy Plasmid Miniprep Kit (Zymo Research) was used to purify plasmid DNA. Sanger-based sequencing reactions using BigDye v3.1 (Applied Biosystems, Foster City, CA) were processed by the University of Wisconsin-Madison Biotechnology Center DNA Sequence Facility.

### Creation of mutants

A fragment of DNA containing *badE* and ∼1.2-kbp flanking DNA up- and downstream of *badE* was PCR amplified, digested with HindIII and BamHI, and ligated into pSUP202 to create pS202badDEF. The *badE* coding region, 350-bp of the 3’ end of *badD*, and 350-bp of the 5’ end of *badF* were deleted from pS202badE by PCR with phosphorylated primers. The resulting PCR product was ligated to an ΩKn^R^ cassette (35) to create pS202ΔbadE. pS202ΔbadE was mobilized into *R. palustris* strains CGA009 and SA008.1.07 via conjugation with *E*. *coli* S17-1. Double crossovers were screened for Kn resistance and ampicillin sensitivity. The presence of the desired *bad* mutations was confirmed by sequencing the appropriate genomic region.

An in-frame, markerless deletion of *hbaB* was constructed using the suicide vector pK18mobsacB (36). Briefly, *hbaB* and ∼0.8-kbp flanking DNA up- and downsteam of *hbaB* was PCR amplified from *R. palustris* genomic DNA, digested with XbaI and HindIII, and ligated into pK18mobsacB to generate pK18hbaB. The *hbaB* coding region was deleted from pK18hbaB by PCR with phosphorylated primers. The resulting PCR product was circularized by ligation to generate pK18ΔhbaB and transformed into *E*. *coli* DH5α. pK18ΔhbaB was introduced into *R. palustris* strains CGA009 and SA008.1.07 by electroporation. Double crossovers were screened for ability to grow on PM-AcY medium with 10% sucrose and Kn sensitivity. The presence of the desired *hbaB* mutation was confirmed by sequencing the appropriate genomic region.

An in-frame, markerless deletion of *vanARB* was constructed in SA008.1.07 using the suicide vector pK18mobsacB (36). Briefly, ∼1.5-kbp up- and downstream flanking regions of *vanARB* were PCR amplified from SA008.1.07 genomic DNA and assembled into pK18mobsacB using the NEBuilder HiFi DNA Assembly Master Mix (New England Biolabs, Ipswich, MA) to create pKΔvanARB. Generation and confirmation of the *vanARB* mutant (SAΔvan) using pKΔvanARB was performed as described above. To generate plasmid pBRvan, a DNA fragment containing the *vanARB* operon was PCR amplified from SA008.1.07 genomic DNA, assembled into the pBBR1MCS-5 vector (37) using the NEBuilder HiFi DNA Assembly Master Mix, and transformed into NEB 5-alpha *E. coli*. After the construction of plasmid pBRvan was confirmed by DNA sequencing, it was introduced into *R. palustris* strains SA008.1.07 and SAΔvan by electroporation. In the same manner, plasmid pBRvanAB was constructed by assembly of pBBR1MCS-5 vector with *vanA* and *vanB* genes amplified from SA008.1.07 genomic DNA, confirmed and transformed to SAΔvan. Transformants were selected on PM-AcY gentamycin plates and confirmed by PCR and DNA sequencing. Gentamycin was added to maintain pBRvan and pBRvanAB.

In-frame, markerless deletions of *rpa2160*, *rpa4286* and *rpa1972* in SA008.1.07, and *rpa1972* in CGA009 were created in the same manner as SAΔvan, creating strains SAΔ2160, SAΔ4286, SAΔ1972, and A9Δ1972, respectively. Primers used for generating gene deletion and expression mutants are shown in Table S2.

## RESULTS AND DISCUSSION

### Isolation of a syringic acid degrading *R. palustris* strain

*R. palustris* CGA009 is reported as being unable to grow photoheterotrophically with syringic acid as the sole organic carbon source (14). To explore the potential for *R. palustris* to evolve the capacity to degrade syringic acid, we established a series of anaerobic cultures in which CGA009 were provided with a combination of syringic acid, benzoic acid, and 4-HBA, the latter two being established growth substrates for this strain (22, 38). Cultures were kept under illumination and anaerobic conditions for at least one week after growth had reached stationary phase. At the conclusion of each growth phase, extracellular samples from each culture were assayed for the presence of aromatic acids. Cultures showing some decrease in extracellular syringic acid concentration were used as an inoculum for new cultures containing an equal or higher proportion of syringic acid in the medium (Figure 1). This process was iterated five times with increases in the proportion of syringic acid in the media, until cells were growing on media in which syringic acid represented 80% of the organic carbon added (measured as COD). The highest performing culture at this stage, as determined by total syringic acid consumption from the media (culture 5.14 in Figure 1), was plated photoheterotrophically onto solid media containing this compound as the sole source of organic carbon, and fourteen colonies were picked after two weeks of incubation. The isolated colonies were then used to inoculate separate liquid photoheterotrophic cultures containing syringic acid as the sole source of organic carbon, the highest performing of which were used as incubations for a second round of liquid photoheterotrophic growth on medium containing syringic acid as the sole organic carbon source. From a second anaerobic plating (from culture 7.07 in Figure 1), twelve colonies were obtained. To further test that these cells acquired the ability to grow solely on syringic acid, cells in isolated colonies were first grown photoheterotrophic on succinate and then subcultured to a medium containing syringic acid as the sole photoheterotrophic carbon source. The isolate that degraded the most syringic acid under photoheterotrophic conditions (Figure 2), hereafter referred to as strain SA008.1.07, was selected for further testing.

**Figure 1.**
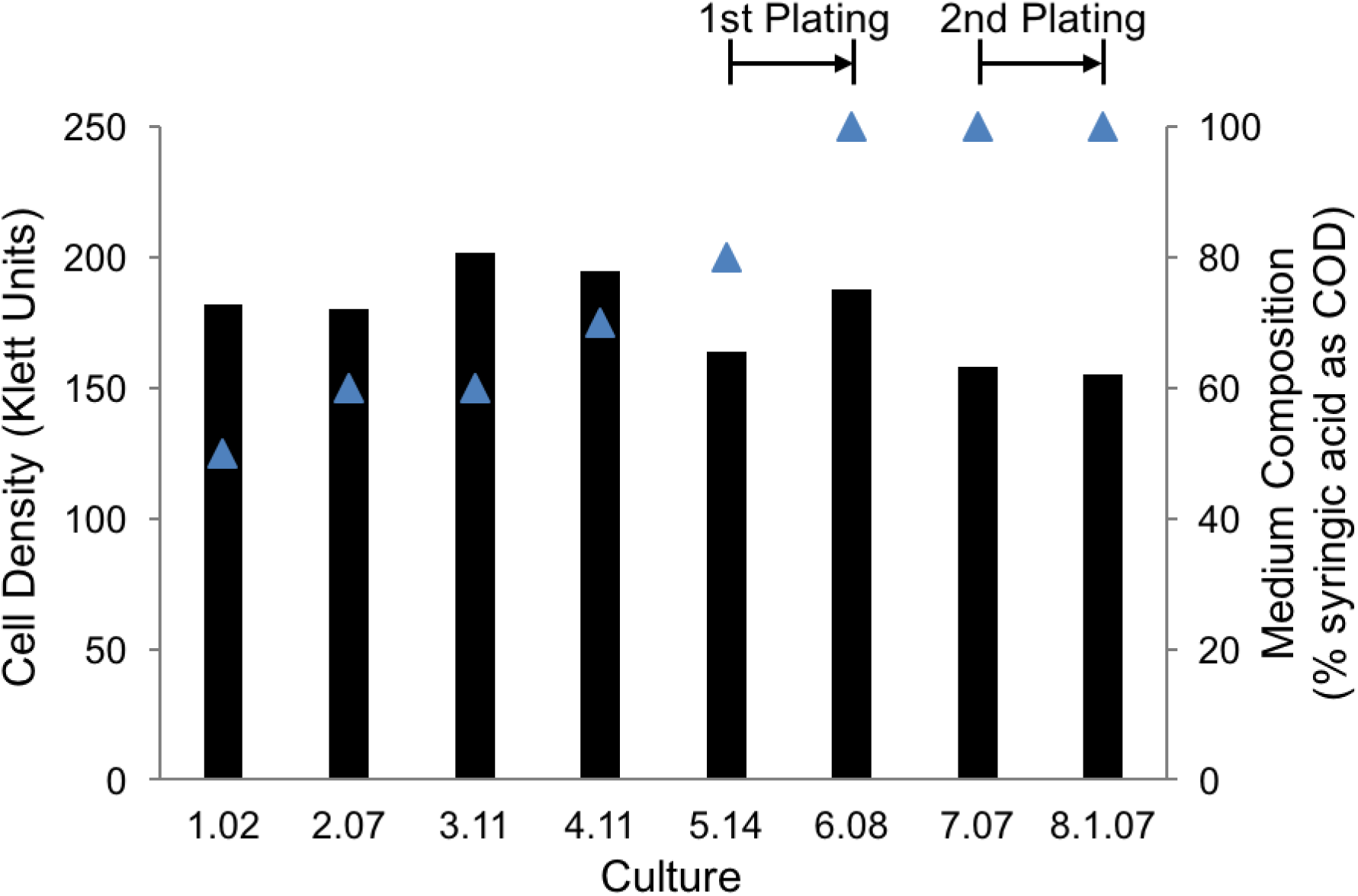
Final cell density (bars, Klett units) and percentage of syringic acid in the culture medium (blue triangles) during sequential anaerobic incubations. Culture 1.02 was started from a colony of *R. palustris* CGA009 that did not exhibit significant metabolism or growth on syringic acid as a sole carbon source. Each culture was seeded from a subculture of the prior one, except in the two instances indicated as 1^st^ plating and 2^nd^ plating in the figure. Cells were plated and single colonies selected for isolation prior to inoculation of cultures 6.08 and 8.1.07. The initial COD of the medium, used as a measurement of bioavailable organic carbon, was maintained at 1g COD/L in all cultures by decreasing the proportion of benzoic acid and 4-HBA upon increases in syringic acid. All cultures were grown anaerobically at 30 °C in sealed glass tubes under constant illumination.

**Figure 2.**
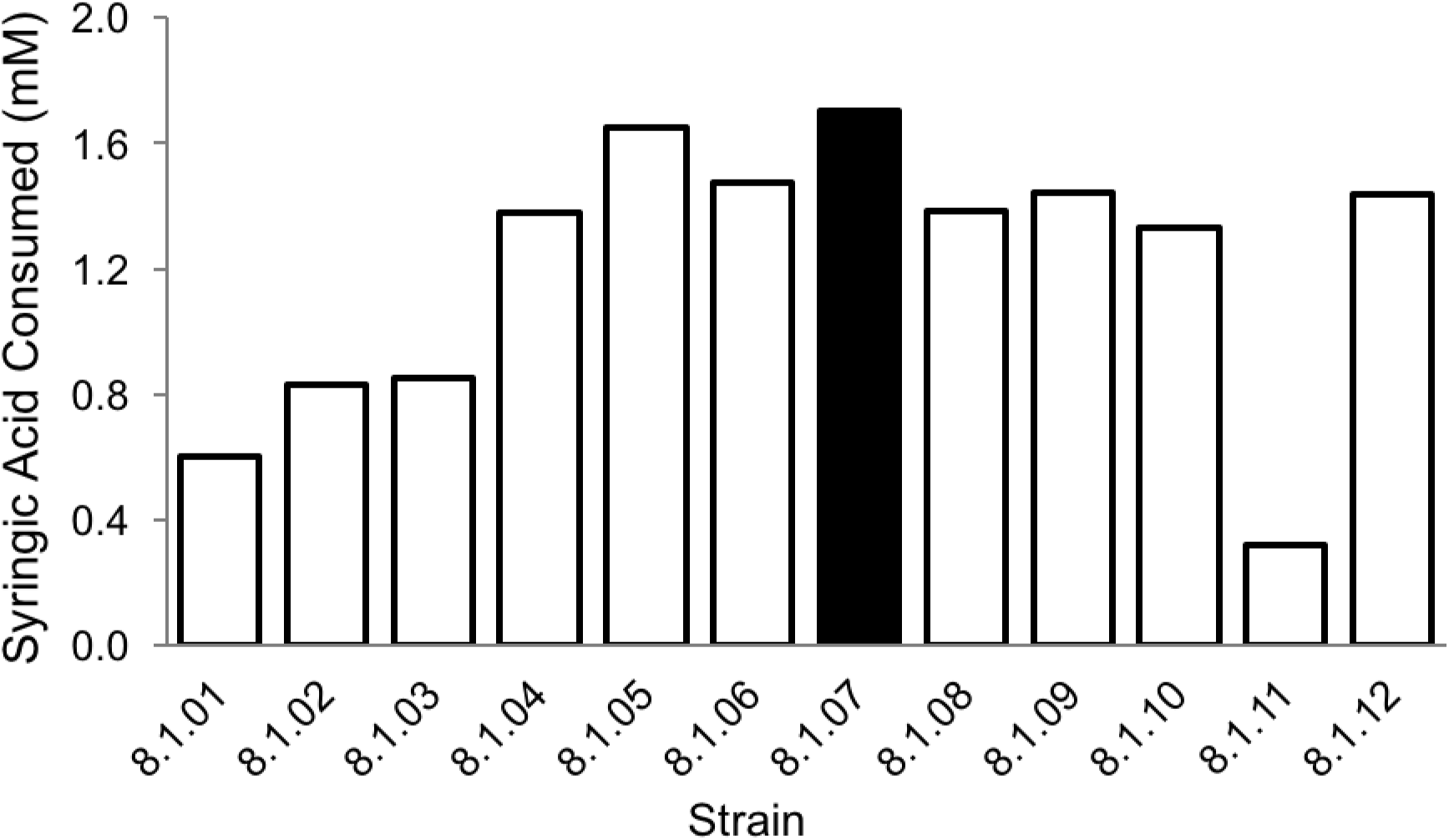
Syringic acid consumption by twelve strains isolated from culture 7.07 (Figure 1). Strain SA008.1.07 had the highest syringic acid transformation and was selected for further study. The initial concentration of syringic acid in these cultures was 3.47 mM.

### Identification of 3,5-dimethoxy-1,4-benzoquinone (DMBQ) as a compound that accumulates extracellularly during growth on syringic acid by SA008.1.07

We found that when SA008.1.07 used syringic acid as a sole source of organic carbon under anaerobic, photoheterotrophic conditions (Figure 3), an orange-yellow tint appeared during early stages of culture growth. However, as growth progressed, the color of the culture became dark and distinguishable from the deep-red color of the accumulating biomass.

**Figure 3.**
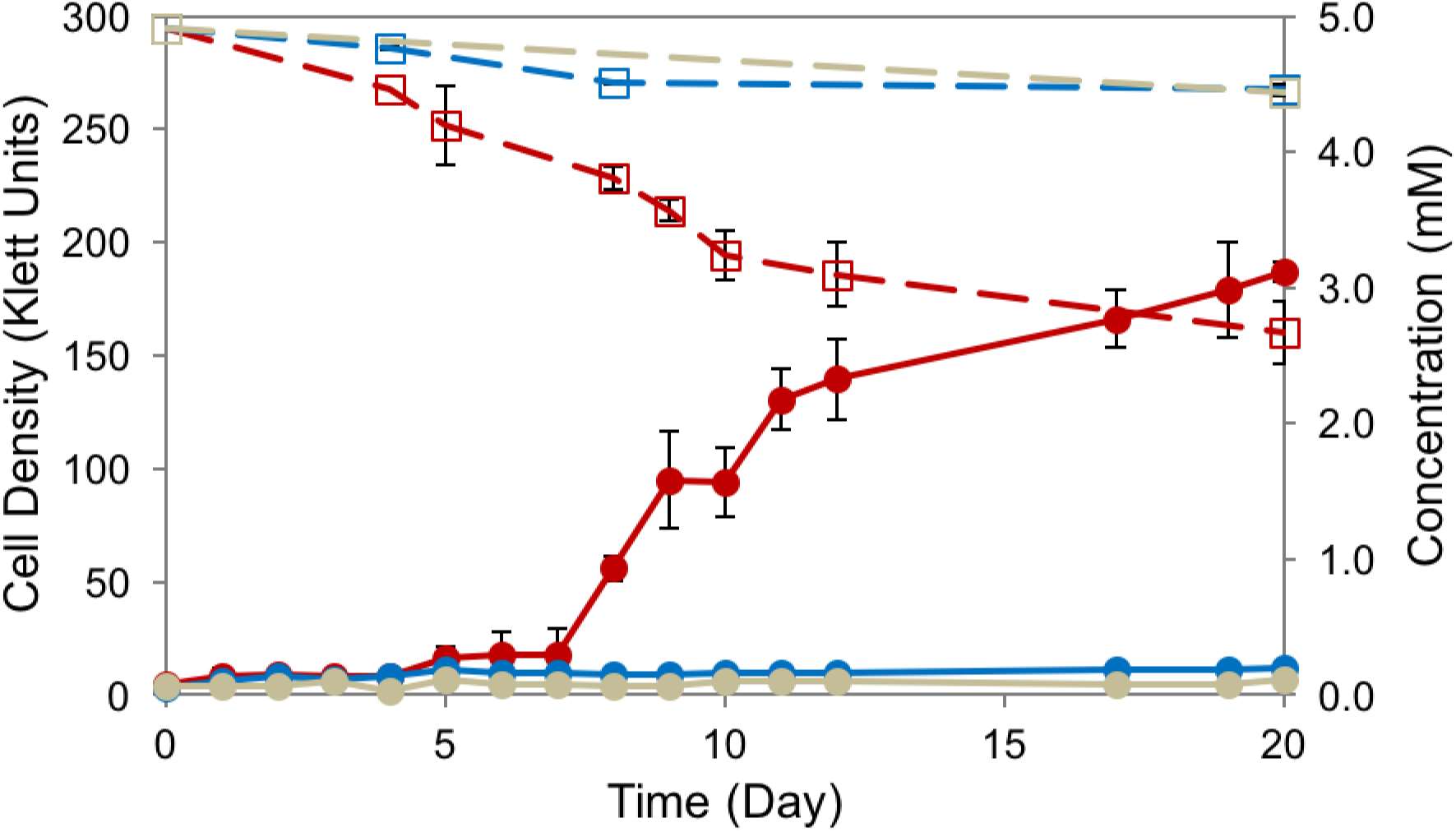
Anaerobic growth of *R. palustris* SA008.1.07 (red) on 5 mM syringic acid, compared to parent strain CGA009 (blue), and light-exposed abiotic control (grey). Solid lines are showing growth in Klett units (●), dashes are tracking concentrations of syringic acid (□). SA008.1.07 consumed approximately half of the syringic acid initially present in the medium, while CGA009 does not grow on syringic acid. Error bars represent standard deviations of experiments performed in triplicate.

HPLC analysis of the media before and after growth of SA008.1.07 revealed the accumulation of a light-absorbing unknown product that eluted at 8.4 min (Figure 4A). By analyzing standards of aromatics that are known or potential syringic-acid degradation by-products (5-hydroxyvanillic acid, vanillic acid, protocatechuic acid) by HPLC, we determined that none of these compounds were found at detectable levels in supernatants from SA008.1.07 cultures. An LC-MS/MS examination of the extracellular unknown indicated an m/z ratio of 169.05 (Figure 4B). For further analysis of this unknown, an extractive procedure was performed on the medium, partitioning the compounds into EtOAc or DCM (See Materials and Methods), and both fractions were analyzed by NMR. Syringic acid was identified as the major product in the ^1^H NMR of the DCM extract, based both on its spectrum and a comparison to that of a commercially purchased standard (Figure 4E). The ^1^H NMR of the EtOAc extract (Figure 4E) contained two major peaks, indicative of methoxy groups and hydrogen atoms on an aromatic ring. Neither of these signals were split, indicating a lack of coupling to adjacent hydrogen atoms in the compound. The predicted molecular weight of the unknown (∼168.05 based on the positive ionization MS spectrum) and the ^1^H NMR pattern suggested that 3,5-dimethoxy-1,4-benzoquinone (DMBQ) was the compound that accumulated during growth on syringic acid. Indeed, NMR (Figure 4E) and MS analysis (Figure 4D) of a commercial DMBQ standard, which also has an orange-yellow tint (CAS number 530-55-2) was indistinguishable from that of the extracellular product that accumulates when SA008.1.07 is grown on syringic acid.

**Figure 4.**
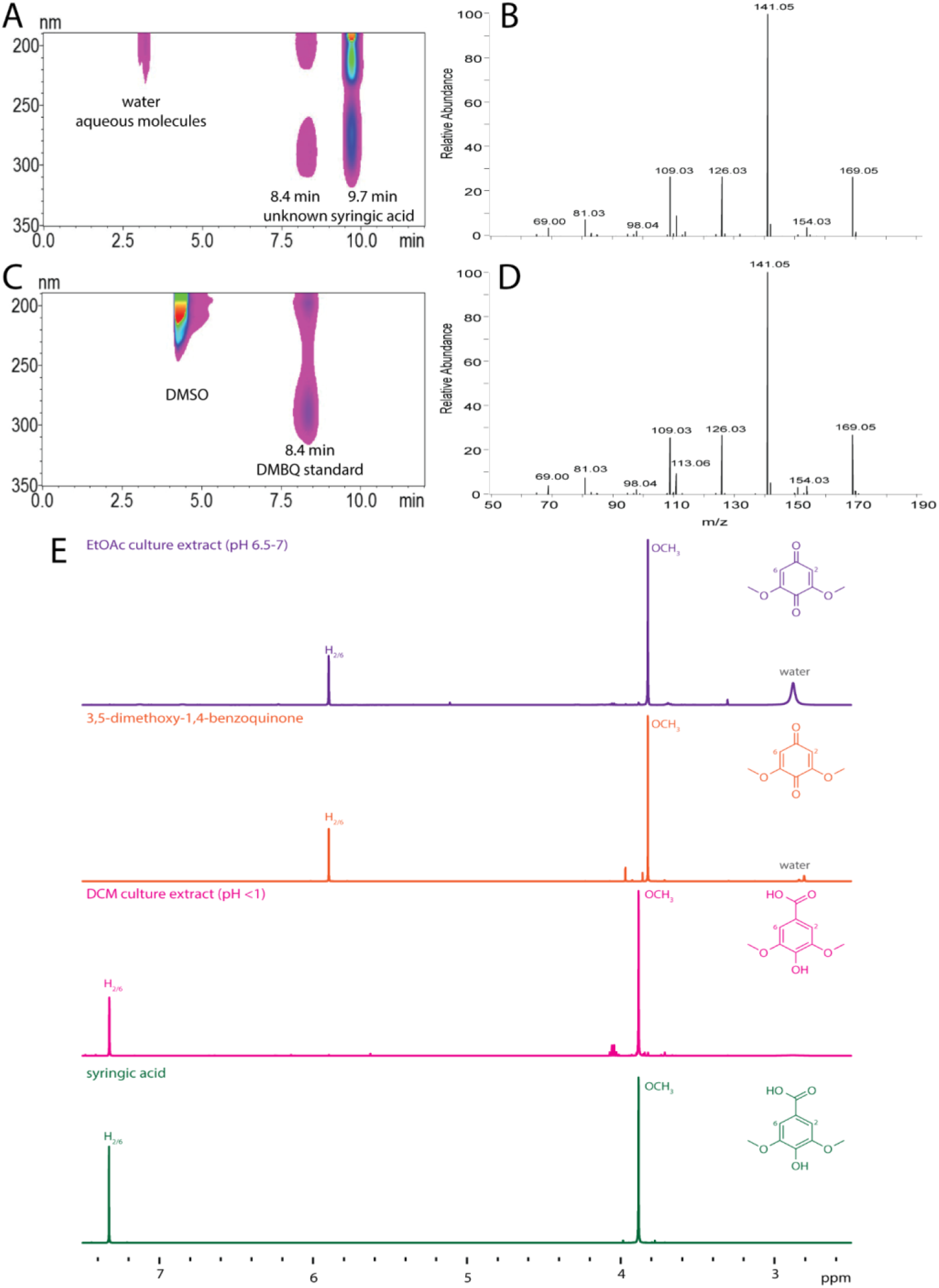
Identification of DMBQ as a soluble extracellular product of SA008.1.07-grown syringic acid cultures. (A) HPLC contour view of PM-syringic acid medium after SA008.1.07 growth, showing peaks at 8.4 and 9.7 minutes, the latter corresponding with syringic acid. (B) LCMS/MS trace of compound isolated from collected peak at 8.4 minutes suggests an m/z ratio of 169.04 g/mol (molecular weight ∼ 168 g/mol). (C) HPLC contour view of DMBQ standard, showing retention time match to the unknown peak in panel A. Peak at 4 minutes is DMSO. (D) LCMS/MS trace of commercially purchased DMBQ showing match to the MS spectrum of unknown peak in panel B. (E) NMR trace of EtOAc extracted culture medium, authentic DMBQ standard, DCM extracted culture medium, and authentic syringic acid standard.

### DMBQ inhibits growth of *R. palustris* SA008.1.07

Since syringic acid was not totally degraded in the SA008.1.07 cultures (Figure 3), we investigated whether the presence of DMBQ affected syringic acid metabolism by this strain. In one test of this hypothesis, we analyzed photoheterotrophic growth of SA008.1.07 in cultures containing 3 mM syringic acid and varying concentrations of DMBQ (Figure 5). When the initial DMBQ concentration was 0.15 mM or above, we observed complete inhibition of growth (as scored by cell density) and of syringic acid degradation (Figure 5). In experiments with initial DMBQ concentrations of less than 0.15 mM, growth and syringic acid degradation occurred, and extracellular DMBQ concentrations increased to about 0.19 mM. Thus, the results of this experiment suggested that, at the range of concentrations tested, DMBQ had an inhibitory effect on syringic acid degradation and cell growth. The inhibitory effect increased as the DMBQ concentration increased, suggesting that the buildup of DMBQ in media containing syringic acid can prevent its total degradation by SA008.1.07. To test this hypothesis, we added 0.3 mM DMBQ (a concentration that approximates the amount found in stationary phase syringic-acid grown cultures) to an SA008.1.07 culture when growth on syringic acid was detected (Figure S1). We found that the addition of 0.3 mM DMBQ arrested growth and blocked further syringic acid degradation in this culture when compared a control not receiving any added DMBQ.

**Figure 5.**
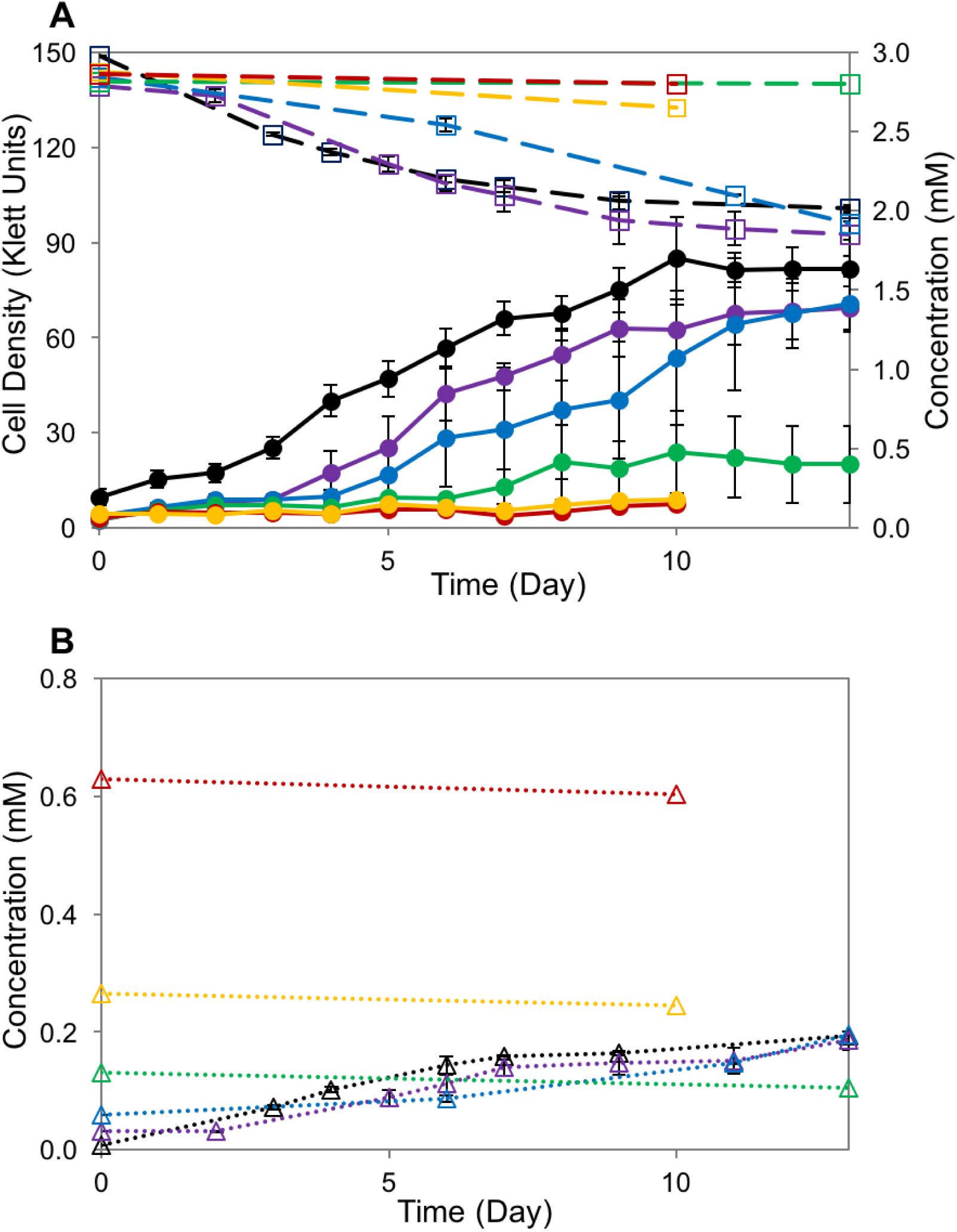
Effect of DMBQ on syringic acid degradation by SA008.1.07. Cultures received 3 mM syringic acid and various starting concentrations of DMBQ (black 0 mM, violet 0.03 mM, blue 0.06 mM, green 0.15 mM, yellow 0.3 mM, red 0.6 mM). (A) Solid lines show cell density in Klett units (●); dashed lines show syringic acid concentration (□). (B) DMBQ concentrations. As the initial concentration of DMBQ increased, cell growth and syringic acid degradation decreased. Cultures with DMBQ concentrations 0.15 mM or greater showed no growth.

To test whether the negative impact of DMBQ on growth was seen in cells grown in the presence of other aromatic substrates, we tested its effects on photoheterotrophic cultures grown on equimolar amounts of benzoic acid and 4-HBA. In this case, we found that addition of 0.3 mM DMBQ to growing SA008.1.07 cultures reduced the rates of growth and of aromatic degradation compared to a control not receiving DMBQ (Figure S2). However, the extracellular DMBQ concentrations decreased in these cultures, suggesting a slow rate of DMBQ degradation that was not evident in experiments with syringic acid. To test the effect of DMBQ on cells growing on a non-aromatic substrate, SA008.1.07 was grown on succinate with varying concentrations of DMBQ (Figure S3). In this case, a lag phase was observed when DMBQ concentrations were 0.06 and 0.3 mM, and complete growth inhibition observed at 0.6 mM. There is also apparent degradation of DMBQ in these cultures (Figure S3). These results indicate that the inhibitory effect of DMBQ on growth or substrate utilization is not specific to cells that are using syringic acid as a sole organic carbon source. However, the inhibitory effect of exogenous DMBQ was more pronounced in cultures growing on aromatic substrates than when using succinate as an organic carbon source. Furthermore, the evidence obtained with these experiments is not sufficient to determine the source of DMBQ. For instance, a benzoquinone has been described as a toxic intermediate in the degradation pathway of pentachlorophenol by *Sphingobium chlorophenolicum* (39). The decrease in DMBQ concentration observed in experiments with 4-HBA and succinate could be a result of either DMBQ being slowly degraded, or reacting with cellular components as described for tetrachlorobenzoquinone in *S. chlorophenolicum* (39).

### Syringic acid degradation by *R. palustris* SA008.1.07 does not require the BAD pathway

To date, the only known route for photoheterotrophic degradation of aromatic compounds in *R. palustris* is through the BAD pathway (19) (Figure S4). To examine the role of the BAD pathway in syringic acid degradation by SA008.1.07, we created SAΔbadE, a mutant of this adapted strain lacking the benzoyl-CoA reductase gene. This deletion is sufficient to block anaerobic degradation of all tested aromatic substrates in wild type *R. palustris* CGA009 (19). We found that the SAΔbadE mutant strain lacks the ability to consume benzoic acid or 4-HBA, as expected (Table 2). However, we also found that SAΔbadE grew on syringic acid, exhibiting a similar behavior to that of the parent strain SA008.1.07 (Figure 6). We also examined the role of the peripheral HBA pathway, responsible for conversion of 4-HBA into benzoyl-CoA (Figure S4), in the growth of strain SA008.1.07 on syringic acid. To do this, we created SAΔhbaB, a mutant of SA008.1.07 which lacks the 4-hydroxybenzoyl-CoA reductase gene that is known to be required for 4-HBA metabolism in *R. palustris* CGA009 (40). As expected, we found that the SAΔhbaB mutant lacks the ability to degrade 4-HBA, yet it can degrade benzoic acid (Table 2). As with the SAΔbadE mutant, we found that SAΔhbaB maintained the ability to grow on and degrade syringic acid (Figure 6).

**Figure 6.**
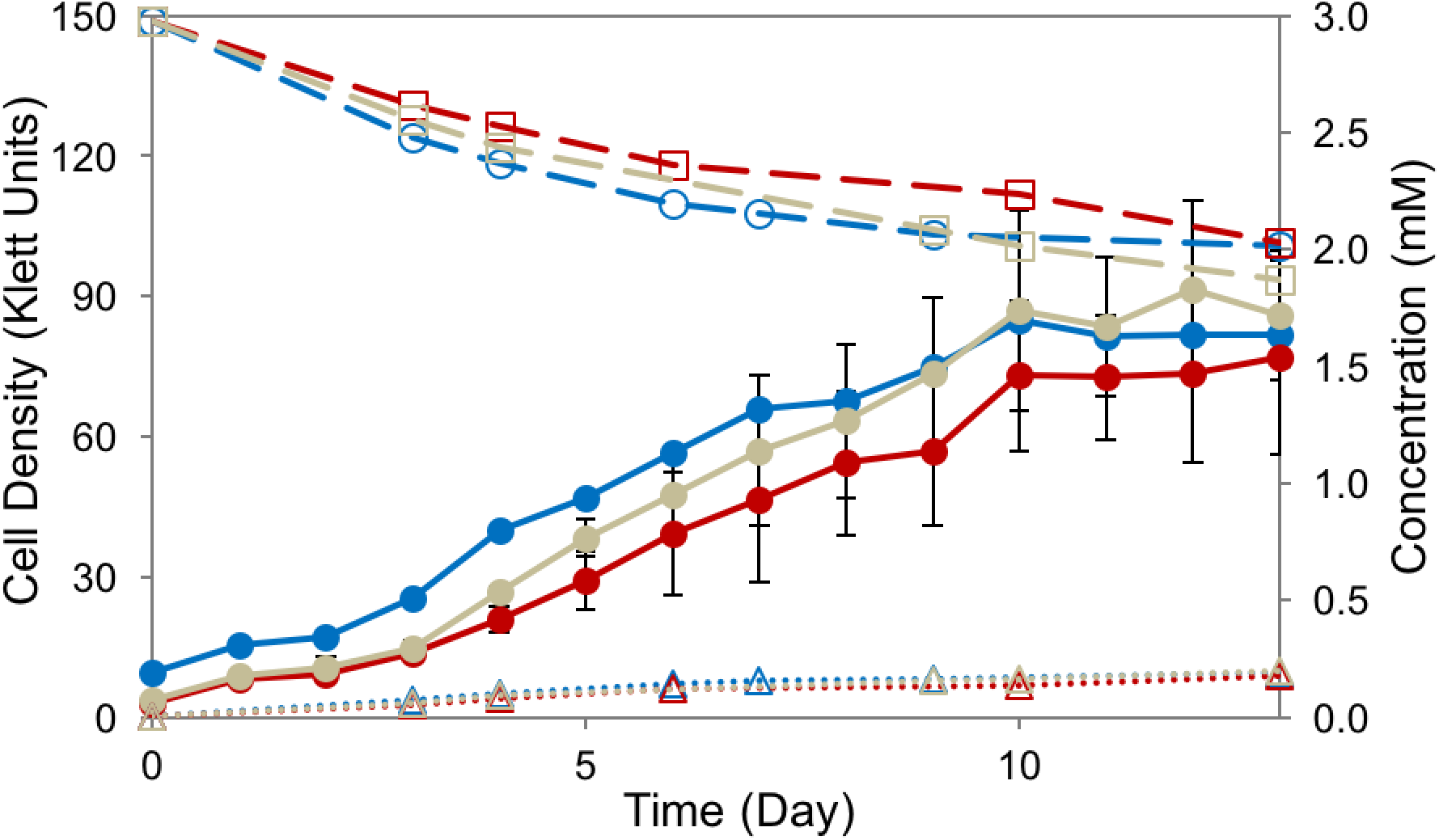
Photoheterotrophic degradation of syringic acid by *R. palustris* strains SA008.1.07 (blue), SAΔbadE (red), and SAΔhbaB (grey). Solid lines are showing cell density in Klett units (●), dashed lines show concentrations of syringic acid (□), and dotted lines show DMBQ concentrations (Δ).

**Table 2.** Endpoint analysis of *R. palustris* SA008.1.07 *bad* and *hba* mutants grown in PM media containing benzoic acid (1.41mM) and 4-HBA (1.63mM).

| **Culture** | **Benzoic Acid**<br>**(mM)** | **4-HBA**<br>**(mM)** | **Final Cell Density**<br>**(Klett Units)** |
| --- | --- | --- | --- |
| SA008.1.07 | ND | ND | 172 |
| SAΔ*badE* | 1.36 | 1.45 | 18 |
| SAΔ*hbaB* | ND | 1.43 | 81 |

ND = not detected

From these experiments, we conclude that neither the peripheral HBA pathway, nor the BAD pathway is required for the degradation of syringic acid by *R. palustris* SA008.1.07. This was a surprising result because the BAD pathway is the only known route for anaerobic aromatic metabolism in *R. palustris* (16, 19).

### Growth on syringic acid does not induce expression of BAD pathways in *R. palustris* SA008.1.07

We used RNA-seq to compare global changes in transcript levels in cultures of SA008.1.07 grown on syringic acid, 4-HBA, and succinate (Table S1). Comparing growth on 4-HBA to growth on succinate revealed the expected increase in transcript abundance of genes involved in the BAD pathway and the peripheral HBA pathway (Table 3). This is consistent with the above finding that SA008.1.07 uses the BAD pathway for 4-HBA metabolism (18). However, the abundance of transcripts from these genes was much lower and mostly not significantly differentially expressed (*p* > 0.05) when comparing growth of SA008.1.07 on syringic acid and succinate (Table 3). Therefore, in addition to SA008.1.07 not needing the BAD and HBA pathways for growth on syringic acid (Figure 6), the transcriptomics data show that growth in the presence of syringic acid does not induce expression of known genes within the BAD and HBA pathways.

**Table 3.** Fold change of genes predicted to be associated with the BAD and peripheral pathways when comparing growth on 4-HBA or syringic acid (SA) to succinate.

| Gene | Name | Predicted Product | Log<sub>2</sub> (Fold Change) |  |
| --- | --- | --- | --- | --- |
|  |  |  | 4-HBA to Succinate | SA to Succinate |
| *rpa0669* | *hbaA* | 4-hydroxybenzoate-CoA ligase | 9.98* | 3.63 |
| *rpa0670* | *hbaB* | 4-hydroxybenzoyl-CoA reductase subunit | 8.40* | 3.01 |
| *rpa0671* | *hbaC* | 4-hydroxybenzoyl-CoA reductase subunit | 8.37* | 2.58 |
| *rpa0653* | *badI* | 2-ketocyclohexanecarboxyl-CoA hydrolase | 7.84* | 2.94* |
| *rpa0658* | *badE* | benzoyl-CoA reductase subunit | 7.68* | 0.36* |
| *rpa0659* | *badF* | benzoyl-CoA reductase subunit | 7.39* | 1.13 |
| *rpa0660* | *badG* | benzoyl-CoA reductase subunit | 7.10* | 0.95 |
| *rpa0656* | *badC* | alcohol dehydrogenase | 6.82* | 0.83 |
| *rpa0654* | *badH* | 2-hydroxycyclohexanecarboxyl-CoA dehydrogenase | 6.70* | 2.00 |
| *rpa0651* | *aliA* | cyclohexanecarboxylate-CoA ligase | 6.38* | 1.00 |
| *rpa0672* | *hbaD* | 4-hydroxybenzoyl-CoA reductase subunit | 6.35* | 0.83 |
| *rpa0657* | *badD* | benzoyl-CoA reductase subunit | 6.09* | -0.40 |
| *rpa0655* | *badR* | benzoate anaerobic degradation transcription regulator | 5.82* | 1.37 |
| *rpa0652* | *aliB* | cyclohexanecarboxyl-CoA dehydrogenase | 5.70* | 0.87 |
| *rpa0650* | *badK* | cyclohex-1-ene-1-carboxyl-CoA hydratase | 5.62* | 0.69 |
| *rpa0667* | *hbaF* | inner-membrane translocator | 5.50* | 0.98 |
| *rpa0662* | *badB* | ferredoxin | 5.01* | 0.55 |
| *rpa0661* | *badA* | benzoate-CoA ligase | 4.91* | 0.77 |
| *rpa0668* | *hbaE* | ABC transporter subunit substrate-binding component | 4.83* | 0.95 |
| *rpa0665* | *hbaH* | ABC transporter ATP-binding protein | 4.67* | 0.50 |
| *rpa0673* | *hbaR* | hydroxybenzoate anaerobic degradation regulatory protein | 4.19* | 0.46* |
| *rpa0666* | *hbaG* | ABC transporter ATP-binding protein | 4.10* | -0.01 |
| *rpa0664* | *badL* | acetyltransferase | 3.64* | 0.04 |
| *rpa3714* | *pimC* | pimeloyl-CoA dehydrogenase large subunit | 3.60* | 0.67 |
| *rpa3713* | *pimD* | pimeloyl-CoA dehydrogenase small subunit | 3.43* | 0.23 |
| *rpa0663* | *badM* | transcriptional regulator BadM | 3.02* | -0.03 |
| *rpa3717* | *pimF* | enoyl-CoA hydratase | 2.63* | 0.45 |
| *rpa3715* | *pimB* | acetyl-CoA acetyltransferase | 2.58* | -0.22 |
| *rpa3716* | *pimA* | AMP-dependent synthetase/ligase | 2.53* | 0.53 |

“\*” indicates statistically significant ( $p < 0.05$ ).

### Identification of a gene cluster required for syringic acid degradation by *R. palustris* SA008.1.07

The global gene expression analysis was also used to identify genes with increased transcript abundance when SA008.1.07 was grown on syringic acid compared to either 4-HBA or succinate (Table 4). Among the transcripts showing the largest increase in abundance are those derived from genes within a putative *vanARB* (*rpa3619-3621*) operon. The *vanARB* genes are annotated as coding for a GntR-family transcriptional regulator (VanR) (41), homologues of which are known or proposed to act as repressors of the *vanAB* genes (42–45). The VanAB proteins are known or predicted subunits of an enzyme (VanAB) with aromatic ring-hydroxylating activity (16, 42). Homologues of VanAB are known or predicted to contain an oxygen-sensitive iron sulfur cluster that catalyzes the oxidation of vanillic acid to protocatechuic acid and formaldehyde in *Bradyrhizobium diazoefficiens (japonicum)* (46) and *Pseudomonas* sp. strain HR199 (47). In addition, a VanAB homologue in a *Streptomyces* strain has the reported ability to demethylate syringic acid as well as other aromatic compounds (48).

**Table 4.** Transcripts with highest increase in abundance when strain SA008.1.07 is grown on syringic acid compared to growth on succinate.

| Gene | Name | Predicted Product | Log<sub>2</sub> (Fold Change) |  |
| --- | --- | --- | --- | --- |
|  |  |  | SA to Succinate | 4-HBA to Succinate |
| *rpa0910* | - | pirin family protein | 9.70 | 1.95 |
| *rpa2160* | - | 3-oxoacyl-ACP reductase | 7.88* | -1.36 |
| *rpa0909* | *wrbA* | NAD(P)H dehydrogenase (quinone) | 6.80* | 0.28 |
| *rpa3619* | *vanA* | aromatic ring-hydroxylating dioxygenase subunit alpha | 6.66* | 0.73 |
| *rpa2717* | - | hypothetical protein | 6.27* | 4.22 |
| *rpa3621* | *vanB* | oxidoreductase | 6.11* | -0.29 |
| *rpa0005* | *hppD* | 4-hydroxyphenylpyruvate dioxygenase | 6.07 | 1.72* |
| *rpa3620* | *vanR* | GntR family transcriptional regulator | 6.01* | -1.34 |
| *rpa4284* | - | polyisoprenoid-binding protein | 5.95* | -0.71 |
| *rpa4222* | - | hypothetical protein | 5.91* | 1.32* |
| *rpa0319* | - | hypothetical protein | 5.90 | 7.58* |
| *rpa3329* | - | hypothetical protein | 5.87 | 3.42 |
| *rpa1475* | - | hypothetical protein | 5.56 | 0.59 |
| *rpa3631* | - | 3-oxoacyl-ACP reductase | 5.52* | -1.20 |
| *rpa4285* | - | malonic semialdehyde reductase | 5.51* | -1.73 |
| *rpa3565* | - | L,D-transpeptidase | 5.39 | -2.17* |
| *rpa3943* | - | ferritin-like domain-containing protein | 5.09* | 0.55* |
| *rpa4394* | - | isocitrate lyase | 5.05* | 6.21* |
| *rpa3308* | - | ferritin-like domain-containing protein | 5.03 | 1.12 |
| *rpa0320* | - | 4-coumaroyl-homoserine lactone synthase | 5.00 | 6.71* |
| *rpa1089* | - | hypothetical protein | 4.99 | 1.32 |
| *rpa0214* | - | hypothetical protein | 4.92* | 0.66* |
| *rpa4220* | - | L,D-transpeptidase | 4.92 | 0.68 |
| *rpa4286* | - | dioxygenase | 4.90* | -2.03 |
| *rpa2895* | - | Hsp20/alpha crystallin family protein | 4.87 | 0.92* |

Shaded lines indicate that the gene was explored in this study.
“\*” indicates statistically significant ( $p < 0.05$ ).

The increased transcript abundance of the *vanARB* genes, which are associated with aerobic degradation of methoxylated aromatic compounds in other bacteria, was unexpected given that the RNA was isolated from cells grown under anaerobic photoheterotrophic conditions. As described in the Materials and Methods section, for RNA-seq experiments cultures were continuously bubbled with N_2_ and CO_2_ to avoid air entering the cultures. For all other experiments, culture tubes were completely filled with medium leaving no headspace, and when samples were withdrawn from the cultures for chemical analyses, the resulting headspace was flushed with argon gas to prevent introducing air into the cultures. These are standard techniques that have been successfully employed to grow anaerobic bacterial cultures and isolate oxygen-sensitive proteins in their active form (24).

We also monitored the abundance of diagnostic transcripts as a reporter for the presence of oxygen in our photoheterotrophic cultures. Analysis of transcript abundance of photoheterotrophically grown cultures shows that there is relatively low abundance of those encoding HemF, an oxygen-dependent coproporphyrinogen oxidase (RPA1514), or subunits of the low affinity enzymes in the aerobic respiratory chain, such as cytochrome *bd* (RPA1319, RPA4452, and RPA4793-4794) or cytochrome *aa_3_* oxidases (RPA1453, RPA4183 and RPA0831-0836) (Table S3). In contrast, transcripts from genes encoding subunits of the high affinity cytochrome *cbb_3_* oxidase (RPA0015-0019), the oxygen-independent coproporphyrinogen oxidase HemN (RPA1666), those needed for anaerobic growth in the light (15, 49), including ones that encode pigment biosynthetic enzymes or pigment-binding proteins of the photosynthetic apparatus (RPA1505-1507, RPA1521-1548, RPA1667-1668, RPA3568), plus others whose induction requires the global anaerobic regulator FixK (RPA1006-1007, RPA1554) (50, 51) are on average ∼32-fold more abundant in the photoheterotrophic cultures than those mentioned above which are associated with growth in the presence of oxygen (p = 0.01, unpaired t-test) (Table S3). This analysis provides independent experimental evidence that the photoheterotrophic cultures used as a source of RNA or for other experiments in this study were anaerobic.

Nevertheless, to further test whether oxygen influences the ability of SA008.1.07 to degrade syringic acid, we performed two additional experiments. First, when we tested SA008.1.07 for aerobic growth on the methoxylated aromatics syringic acid and vanillic acid (Figure S5), we found that this adapted strain cannot grow on syringic acid aerobically. We also performed growth experiments in which additional steps were taken to eliminate oxygen from the media. For this, we used 100 mL serum bottles, added PM media containing syringic acid, and sealed them with rubber septa and aluminum crimp caps. We then flushed the PM media with argon gas for 20 min, then applied vacuum to remove gases from the bottles, and re-flushed them with argon. This process was repeated three times to remove as much oxygen as possible. As a control that simulated conditions used in the experiments described earlier, another group of 100 mL serum bottles was used, but in this case the bottles were sealed without using the degassing procedure. SA008.1.07 was inoculated into both sets of bottles through sterilized syringes and needles. In these experiments, we observed no significant difference on the growth of SA008.1.07, the consumption of syringic acid, or the production of DMBQ between the degassed bottles and the non-degassed controls (Figure S6), demonstrating that any traces of oxygen potentially present at the initiation of the incubations did not influence the ability of *R. palustris* SA008.1.07 to grow on syringic acid under anaerobic photoheterotrophic conditions.

Based on these results, we proceeded to investigate whether the *vanARB* operon participated in anaerobic syringic acid degradation by SA008.1.07. To do this, we deleted the entire *vanARB* operon in SA008.1.07 (SAΔvan, Table 1), and found that this strain lost its ability to grow anaerobically on syringic acid (Figure 7A). In addition, we found that transforming SAΔvan with a plasmid carrying either the wild-type *vanARB* operon or only wild-type *vanAB* (SAΔvan.pBRvanARB or SAΔvan.pBRvanAB, Table 1) under control of a constitutive promoter rescues the ability of SAΔvan to grown on and degrade syringic acid under anaerobic conditions, although cell densities were lower than in SA008.1.07 (Figure 7A). Thus, we conclude that *vanAB* genes in the *R. palustris van* cluster are required for anaerobic degradation of syringic acid by SA008.1.07. In control experiments, we found that, as expected, the activity of the BAD and HBA aromatic pathways were not affected by the loss of *vanARB*, as SAΔvan is able to grow photoheterotrophically on 4-HBA or benzoic acid (Figure 7B). Placing the same *vanARB* plasmid in the wild-type CGA009 strain (A9pBRvanARB, Table 1) does not confer this strain with the ability to grow on syringic acid (Figure 7A), indicating that yet to be identified genes outside this operon are required for syringic acid metabolism by SA008.1.07.

**Figure 7.**
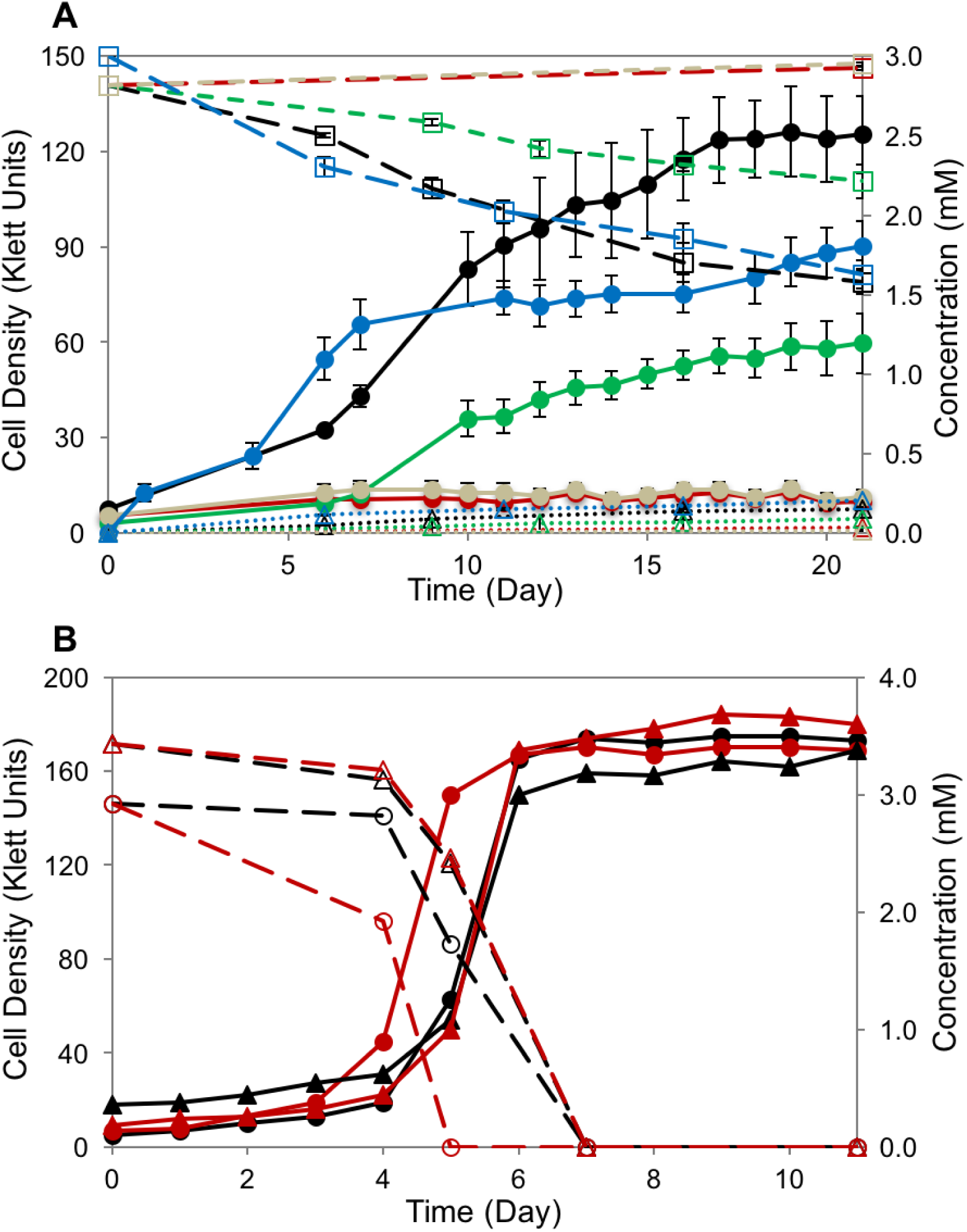
(A) *R. palustris* SAΔvan (red), a mutant culture of SA008.1.07 (black) with the *vanARB* operon deleted, does not grow on syringic acid. The complementation of *vanARB* on expression plasmid pBRVanARB and pBRVanAB restores syringic acid degrading activity in SAΔvan.pBRvanARB (green) and SAΔvan.pBRvanAB (blue). The expression plasmid pBRVanARB does not impart syringic acid degrading activity when inserted into wild-type strain CGA009 (A9pBRvanARB, grey). Solid lines are showing growth in Klett units (●), dashes tracking concentrations of syringic acid (□), and dotted lines tracking DMBQ concentration (Δ). (B) *R. palustris* SA008.1.07 (black) and SAΔvan (red) both grow on benzoic acid (circles) and 4-HBA (triangles). Solid lines are showing growth in Klett units and dashes are indicating aromatic concentrations.

In addition to the genes in the *vanARB* operon, we also tested the effect of deleting two other genes that showed increased transcript abundance during growth on syringic acid. One gene was an oxidoreductase that had one of the highest increases in transcript abundance (*rpa2160*) and the other gene was a dioxygenase with a lower increase in transcript abundance (*rpa4286*) (Table 4). Experiments with deletion mutants of SA008.1.07 lacking these genes, SAΔ2160 and SAΔ4286 respectively (Table 1), showed that neither deletion affected photoheterotrophic growth on syringic acid (Figure S7), indicating that these genes are not required for the breakdown of syringic acid by SA008.1.07.

### Identification of mutations in strains adapted to grow on syringic acid

In an attempt to identify additional mutations that could confer *R. palustris* SA008.1.07 with the ability to grow photoheterotrophically on syringic acid, we re-sequenced strain SA008.1.07 along with 16 other *R. palustris* isolates that had acquired the same metabolic ability by following the same enrichment and isolation experiments (Table S4). When the genome sequences of this panel of isolates were compared to that of *R. palustris* CGA009 (Table S5), only 4 mutations were found in the majority of the strains (Table 5). One mutation was an indel upstream of *rpa0746*, a gene annotated as encoding a *c*-type cytochrome of unknown function. A second mutation was a frameshift in *rpa1972*, a gene annotated as encoding a two-component sensor histidine kinase, for which no function is known. The other two mutations were non-synonymous, causing amino acid changes in *rpa2457*, a hypothetical protein, and *rpa3268*, which encodes the β-subunit of RNA polymerase. No mutations were detected in the *vanARB* operon in any of the syringic acid-metabolizing strains that were sequenced.

**Table 5.** Mutations identified in more than half of the 17 adapted *R. palustris* strains conferring the ability of syringic acid degradation compared with the genome of CGA009.

| Position | Reference | Alteration | Mutation Type | Amino Acid Change | Gene | Name | Function | Occurrence of Mutation |
| --- | --- | --- | --- | --- | --- | --- | --- | --- |
| 826195 | C | A | INDEL | - | Upstream<br>*rpa0746* | - | - | 17 of 17 strains |
| 2221051 | TC | T | Frame Shift | - | *rpa1972* | - | two-component<br>sensor histidine<br>kinase | 17 of 17 strains |
| 2795227 | G | A | Non-Synonymous | Glycine →<br>Aspartic Acid | *rpa2457* | - | hypothetical protein | 16 of 17 strains |
| 3685350 | T | G | Non-Synonymous | Threonine →<br>Proline | *rpa3268* | *rpoB* | RNAP Beta | 11 of 17 strains |

We were unsuccessful in our attempts to delete *rpa2457* and *rpa3268* from SA008.1.07 using the methods used in this study, which is not surprising since both of these genes have been shown to be essential for the growth of *R. palustris* (15). We successfully deleted *rpa1972* in both CGA009 and SA008.1.07, creating strains SAΔ1972 and A9Δ1972, respectively. To test the hypothesis that the observed frameshift in *rpa1972* altered the function of this predicted histidine kinase and somehow influenced syringic acid degradation by SA008.1.07, we evaluated photoheterotrophic growth of both SAΔ1972 and A9Δ1972 on syringic acid. This experiment showed that deletion of *rpa1972* in CGA009 did not enable A9Δ1972 to grow on syringic acid, nor did deletion of this gene in SA008.1.07 prevent SAΔ1972 from growing on syringic acid (Figure S8). Therefore, additional efforts are needed to identify single or synergistic combinations of mutations in SA008.1.07 or other adapted strains that contribute to anaerobic growth on syringic acid.

### Concluding remarks

Meta-methoxylated aromatics are present at significant levels in the lignin of different plants and are potential sources of compounds for industrial applications. In this work, we isolated a strain of *R. palustris* that acquired the ability to use syringic acid as a growth substrate under photoheterotrophic conditions. Our strategy of incrementally exposing cultures to higher concentrations of syringic acid, while at the same time reducing the availability of the known growth substrates benzoic acid and 4-HBA, has been shown to be conducive to adaptation and acquisition of new metabolic activities in *R. palustris* (52, 53) and other bacteria (54). Our analysis of this adapted strain, SA008.1.07 has provided important new knowledge on the bacterial metabolism of syringic acid. First, we found that syringic acid degradation does not occur through or induce expression of the genes in the well-characterized BAD pathway. This finding makes syringic acid the first aromatic compound whose photoheterotrophic metabolism does not utilize the BAD pathway in *R. palustris*. In addition, the increased abundance of *vanARB* transcripts in SA008.1.07 cultures grown in the presence of syringic acid, and the requirement of *vanAB* for growth of this adapted strain on this methylated aromatic provide evidence for a heretofore unknown role of this enzyme in anaerobic metabolism of this compound. Since the previously reported function of *vanAB* is in the aerobic demethylation of vanillic acid (16, 42), our observations suggest that the VanAB enzyme may have an additional unrealized function under anaerobic conditions. Known homologues of VanAB are reported to contain an oxygen-sensitive iron sulfur cluster (47) so our findings reinforce that additional experiments are needed to test the role of this enzyme in anaerobic metabolism of syringic acid.

Our analysis of syringic acid metabolism by *R. palustris* SA008.1.07 sets the stage for further studies of metabolism of this and other aromatics by this and other bacteria, and for evaluating previously unexplored functions of the VanAB enzyme. Elucidating such novel pathways and metabolic functions could expand our ability to use microbial transformations of lignin and other renewable resources as bio-based sources of compounds with potential uses in the energy, chemical, pharmaceutical, and other industries.

## ACKNOWLEDGMENTS

This work was supported by the Department of Energy Office of Science’s Great Lakes Bioenergy Research Center, Grants DE-FC02-07ER64494 and DE-SC0018409, and the National Science Foundation, Grant CBET-1506820. J. Zachary Oshlag was supported by the National Institute of General Medical Sciences of the National Institutes of Health under Award Number T32GM008349. The genome sequencing work conducted by the U.S. Department of Energy Joint Genome Institute, a DOE Office of Science User Facility, is supported by the Office of Science of the U.S. Department of Energy under Contract No. DE-AC02-05CH11231. We thank Daniel L. Gall and Alan Higbee for preliminary work in the identification of DMBQ.

## SUPPLEMENTARY FIGURES

**Figure S1.**
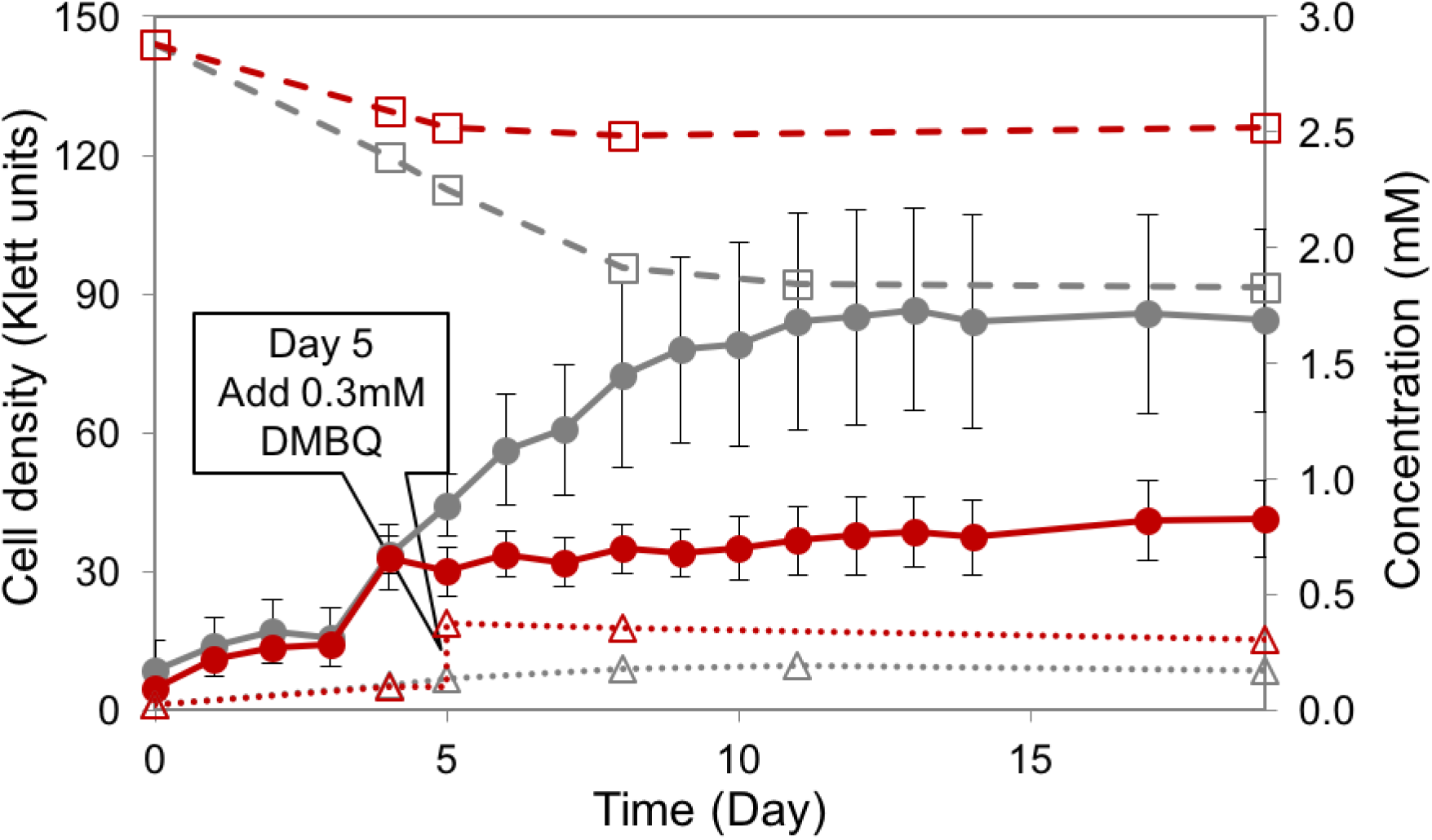
Effect of DMBQ addition on syringic acid degradation by *R. palustris* SA008.1.07. Solid lines show cell density (Klett units), dashes lines show syringic acid concentration, dotted lines show DMBQ concentration. Red lines indicate results for the DMBQ-containing culture and grey lines show results for a control culture not receiving DMBQ. DMBQ (0.3 mM) was added to the culture on Day 5. For this experiment, DMBQ was dissolved in DMSO, and the control culture was provided with DMSO only. Error bars represent standard deviation of experiments performed in triplicate.

**Figure S2.**
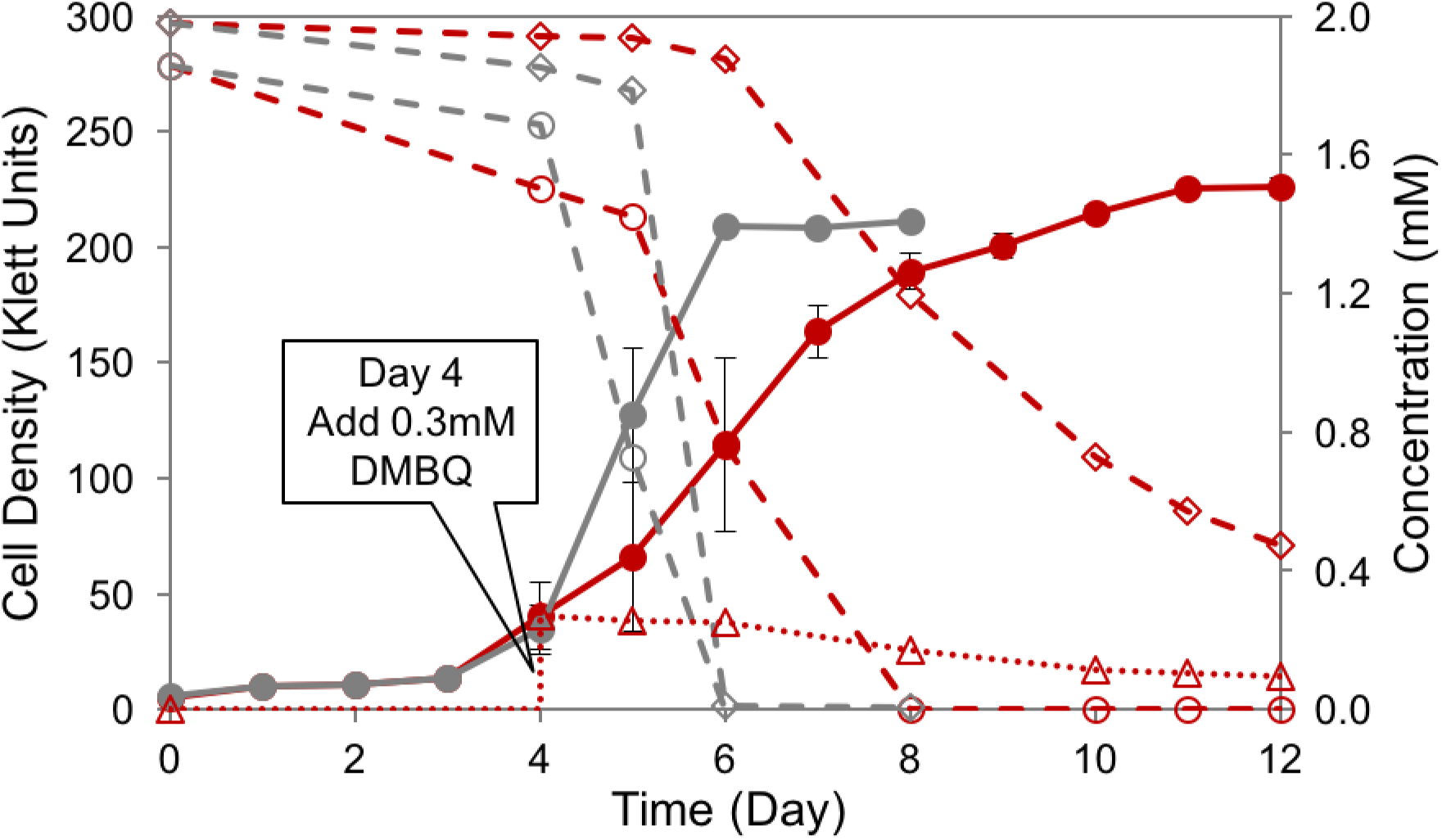
Effect of DMBQ addition on *R. palustris* SA008.1.07 growing on an equimolar amount of benzoic acid and 4-HBA (Initial concentration was 2 mM for each aromatic substrate). Solid lines are showing growth in Klett units (●), dashes tracking concentrations of benzoic acid (○), 4-HBA (◊), and dotted lines tracking DMBQ concentration (Δ). Red lines indicate results for the DMBQ-containing culture. DMBQ (0.3 mM) was added to the culture on Day 4. For this experiment, DMBQ was dissolved in DMSO. Parallel control cultures (in grey) received DMSO without DMBQ.

**Figure S3.**
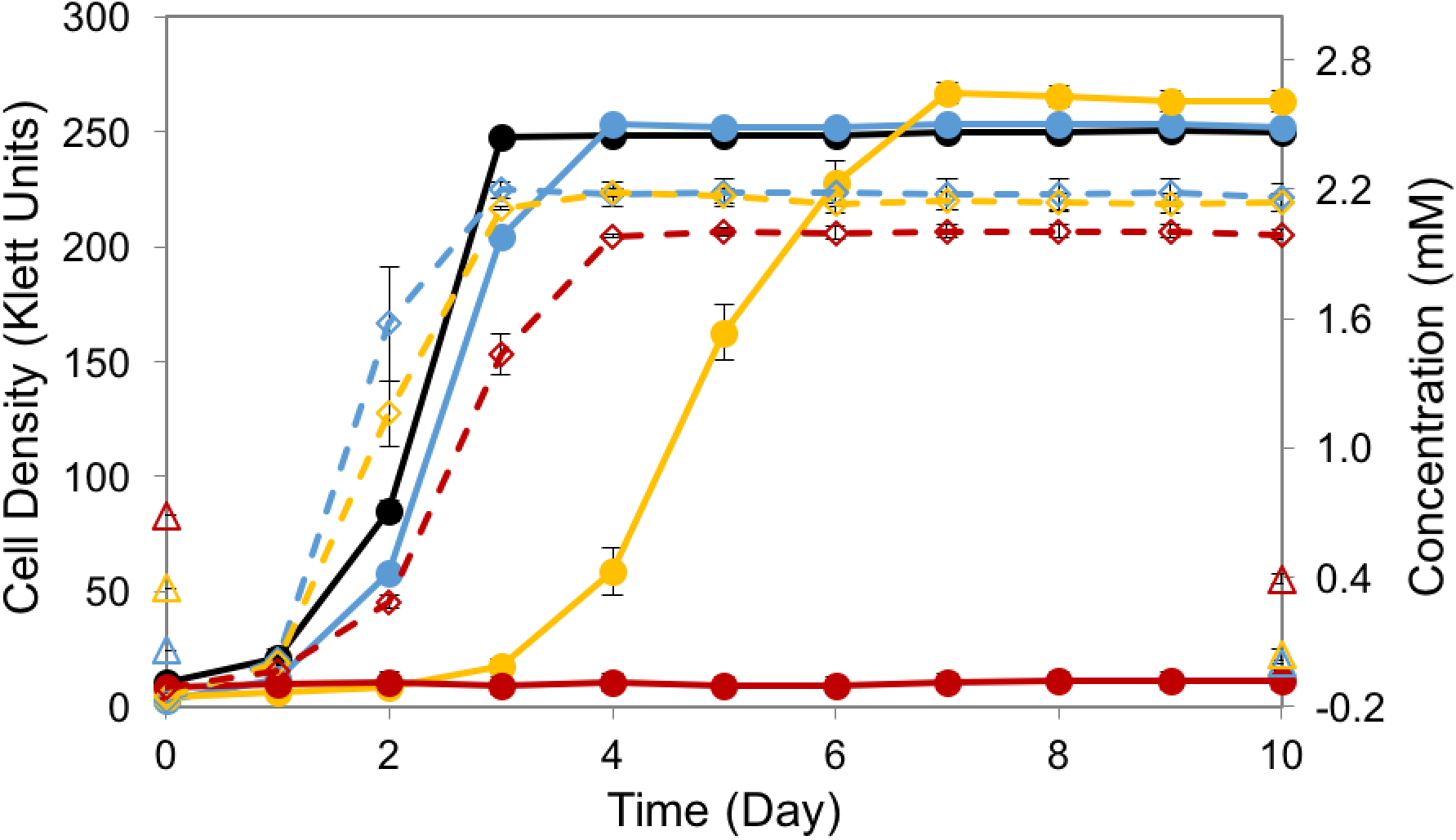
Effect of DMBQ on SA008.1.07 cultures grown on succinate. Solid lines show cell density of cultures received 10 mM succinate and various starting concentrations of DMBQ (black 0 mM, blue 0.06 mM, yellow 0.3 mM, red 0.6 mM). For these experiments, DMBQ was dissolved in DMSO. Dashed lines are showing growth of control cultures received corresponding amount of DMSO without DMBQ. Triangles (Δ) denote concentrations of DMBQ at the beginning and end of the experiment.

**Figure S4.**
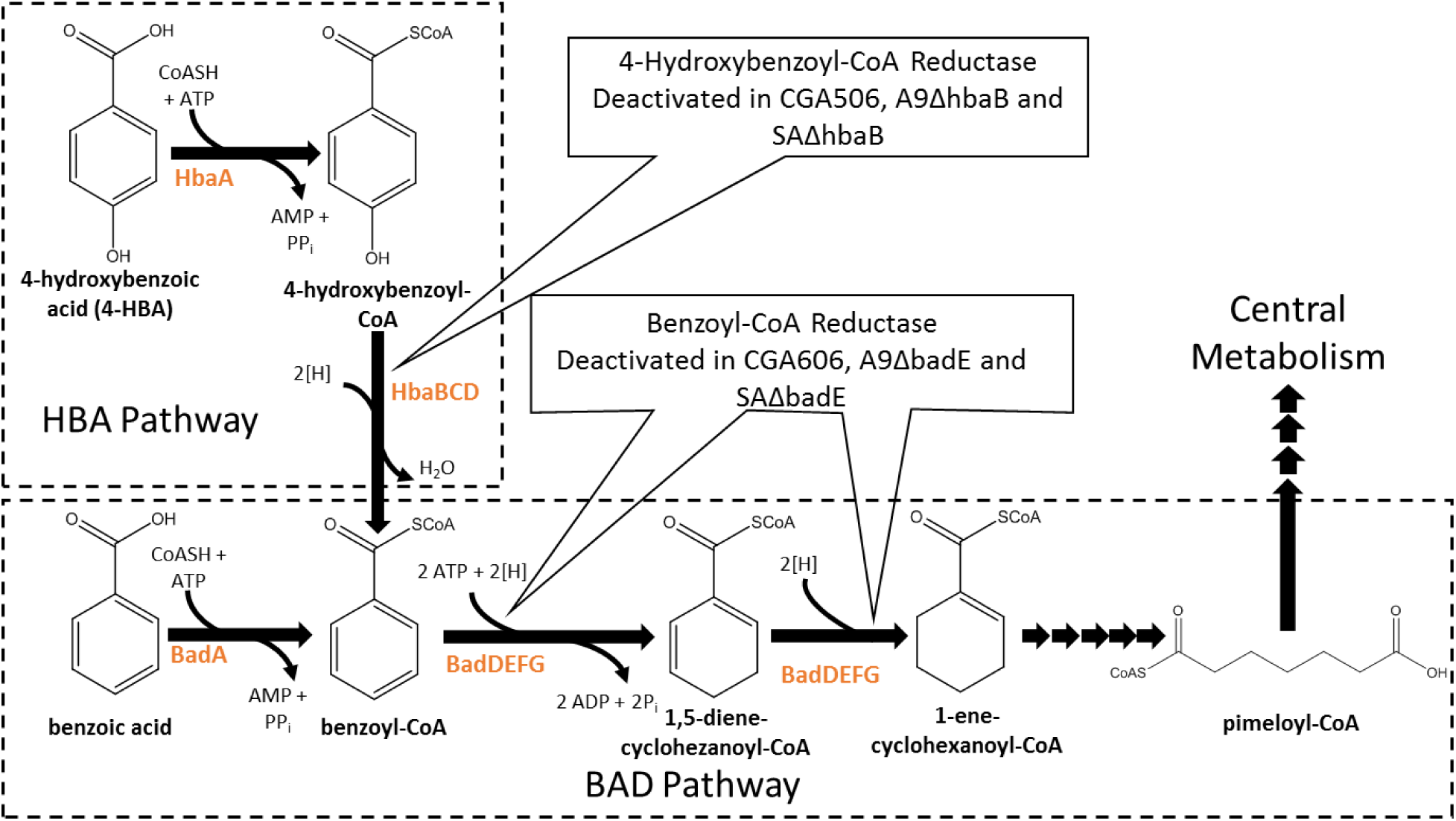
4-Hydroxybenzoic acid (HBA) and benzoic acid degradation (BAD) pathways. These are the only previously established routes for anaerobic degradation of aromatic acids by *R. palustris.* HbaBCD and BadDEFG are oxygen sensitive enzymes.

**Figure S5.**
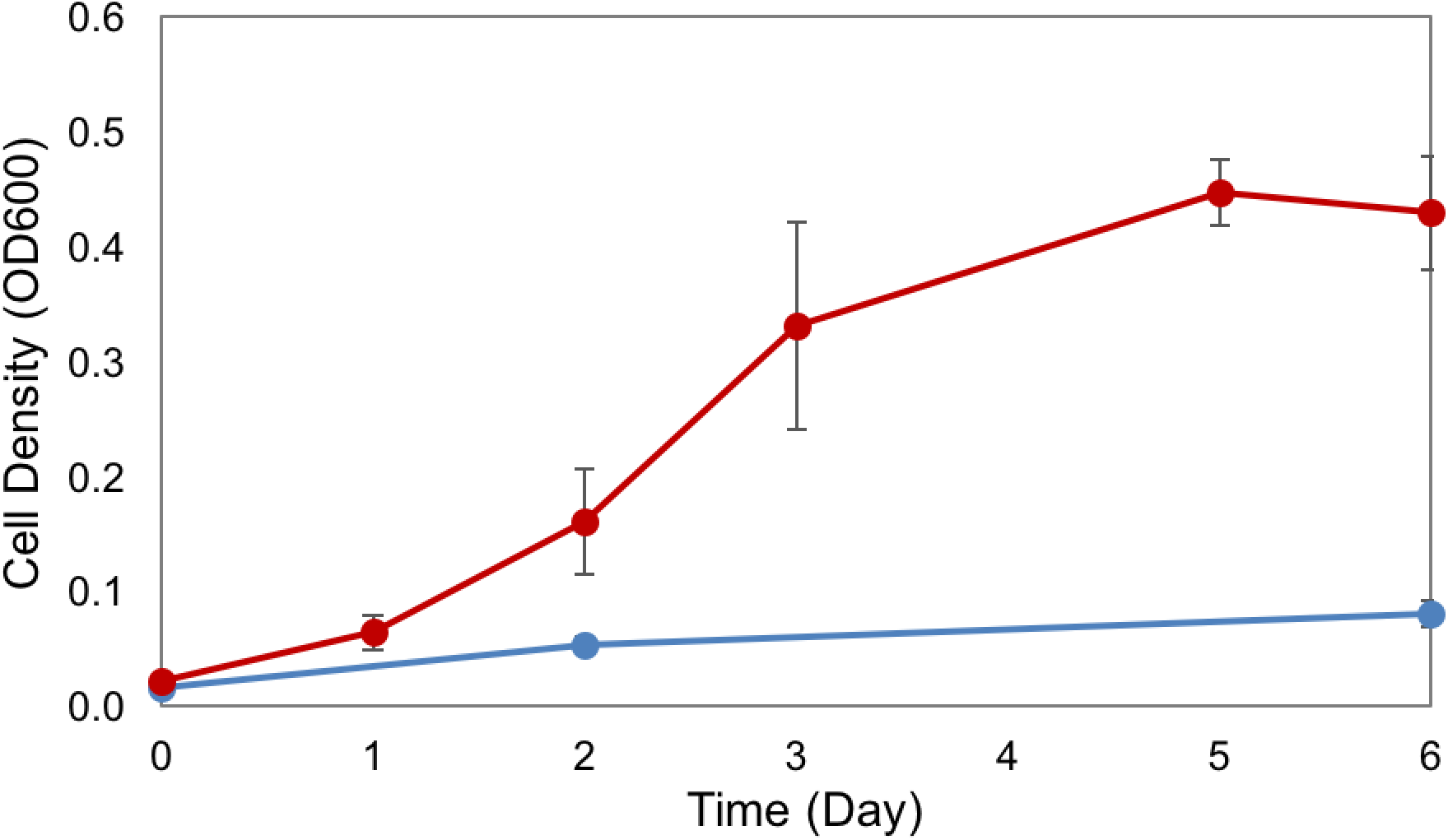
Aerobic growth of SA008.1.07 in 3 mM syringic acid (blue line) or vanillic acid (red line). At the end of experiment, syringic acid was not consumed, while vanillic acid was completely consumed.

**Figure S6.**
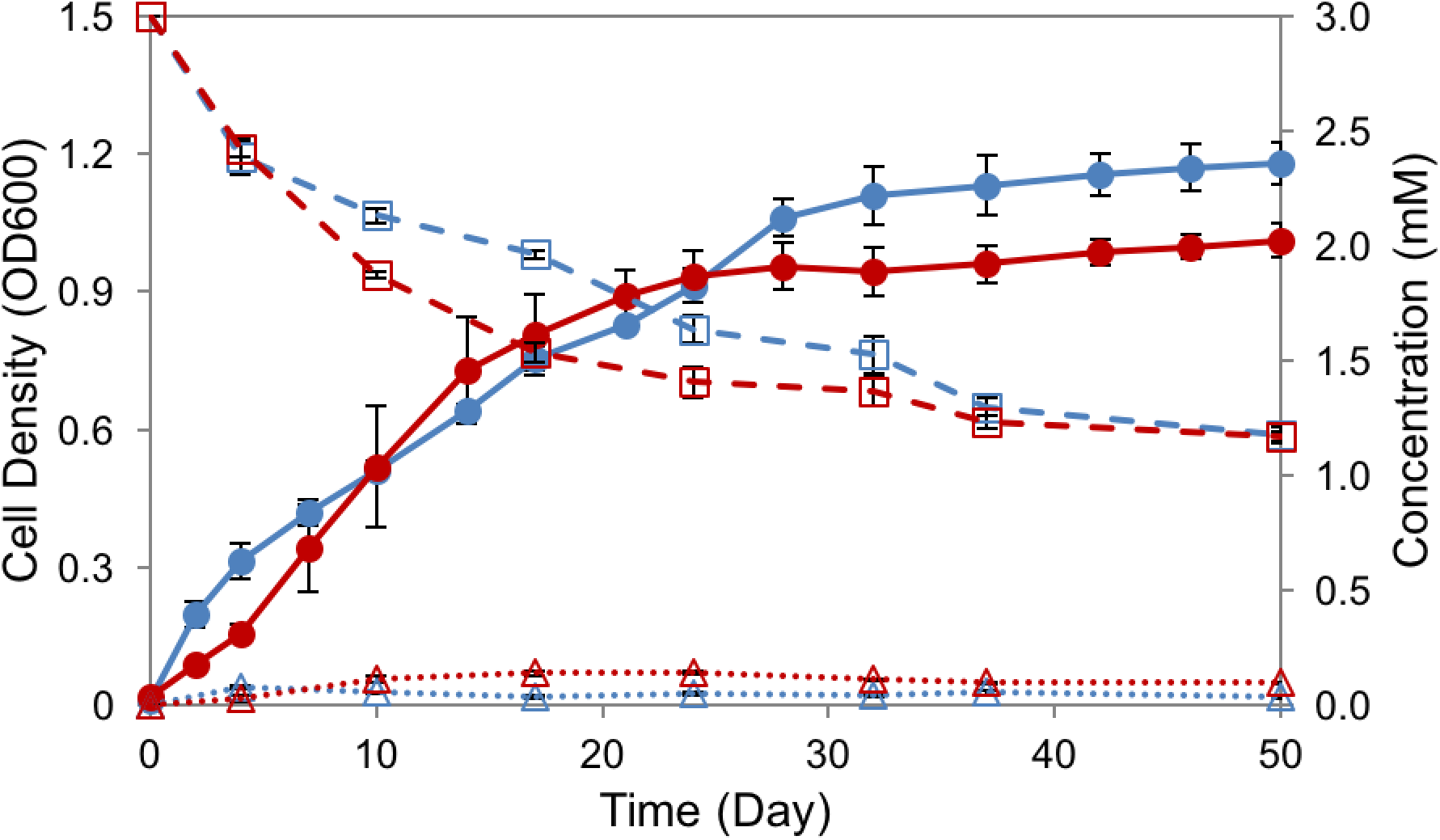
Cultures of SA008.1.07 in 3 mM syringic acid, grown on degassed (blue) and non-degassed (red) serum bottles. Solid lines are showing cell density in OD600 (●), dashes tracking concentrations of syringic acid (□), and dotted lines tracking DMBQ concentration (Δ).

**Figure S7.**
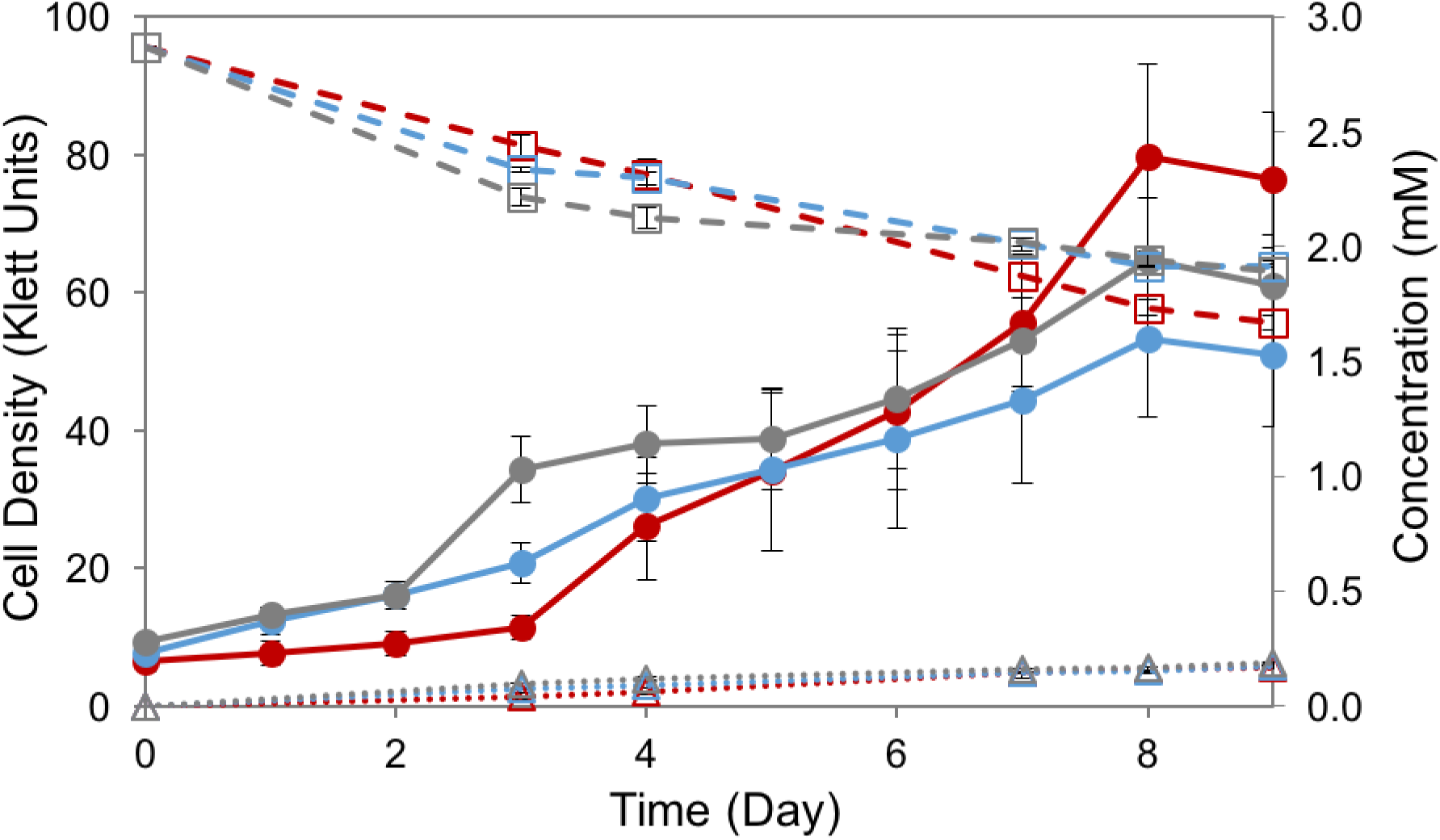
Cultures of *R. palustris* in 3 mM syringic acid. Growth and syringic acid consumption phenotype of SA008.1.07 (red) matches that of deletion strains SAΔ2160 (blue) and SAΔ4286 (grey). The genes that were deleted in these strains *rpa2160* and *rpa4286* do not appear to be necessary for growth of SA008.1.07 on syringic acid. Solid lines are showing growth in Klett units (●), dashes tracking concentrations of syringic acid (□), and dotted lines tracking DMBQ concentration (Δ).

**Figure S8.**
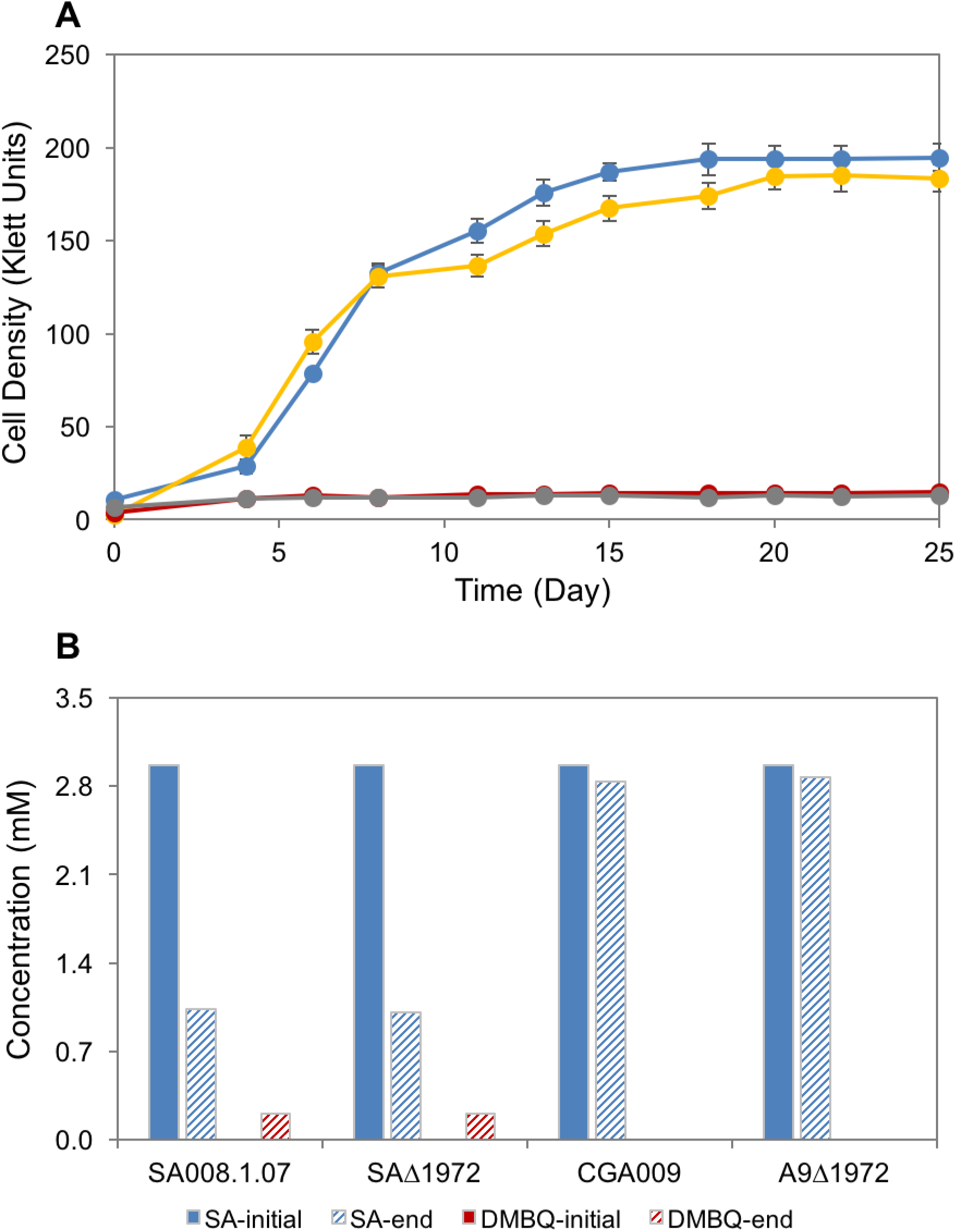
Cultures of *R. palustris* in 3 mM syringic acid. (A) Growth and syringic acid consumption phenotype of SA008.1.07 (blue) and CGA009 (red) matches that of deletion strains SAΔ1972 (yellow) and A9Δ1972 (grey), respectively. (B) Concentration of syringic acid (SA, blue bars) and DMBQ (red bars) in the initial and end-point of the cultures.

